# Molecular Landscape of Anti-Drug Antibodies Reveals the Mechanism of the Immune Response Following Treatment with TNFα Antagonists

**DOI:** 10.1101/509489

**Authors:** Anna Vaisman-Mentesh, Shai Rosenstein, Miri Yavzori, Yael Dror, Ella Fudim, Bella Ungar, Uri Kopylov, Orit Picard, Aya Kigel, Shomron Ben-Horin, Itai Benhar, Yariv Wine

## Abstract

Drugs formulated from monoclonal antibodies (mAbs) are clinically effective in various diseases. Repeated administration of mAbs, however, elicits an immune response in the form of anti-drug-antibodies (ADA), thereby reducing the drug’s efficacy. Notwithstanding their importance, the molecular landscape of ADA and the mechanisms involved in their formation are not fully understood. Using a newly developed quantitative bio-immunoassay, we found that ADA concentrations specific to TNFα antagonists can exceed extreme concentrations of 1 mg/ml with a wide range of neutralization capacity. Our data further suggest a preferential use of the λ light chain in a subset of neutralizing ADA. Moreover, we show that administration of TNFα antagonists result in a vaccine-like response whereby ADA formation is governed by the extrafollicular T cell-independent immune response. Our bio-immunoassay coupled with insights on the nature of the immune response can be leveraged to improve mAb immunogenicity assessment and facilitate improvement in therapeutic intervention strategies.

## Introduction

More than 30 years since the approval of the first therapeutic monoclonal antibody (mAb) for clinical use, the therapeutic mAb market has expanded exponentially, establishing mAbs as one of the leading biopharmaceutical therapeutic modalities (1). Although mAbs hold significant promise for improving human health, their repeated administration is often highly immunogenic and can elicit an undesirable anti-drug antibody (ADA) response (2). The formation of an ADA response interferes with the effect of the drug or neutralizes it thereby altering the drug’s pharmacokinetic (PK) and pharmacodynamic (PD) properties and reducing its efficacy (3), and eventually may lead to a severe adverse immune reaction in humans (4).

Immunogenicity of mAbs and the formation of an ADA response has been suggested to be dependent on the interplay between factors related to the drug itself (e.g., non-human sequence, glycosylation, impurities, aggregation), to the patient (e.g., disease type, genetic factors, concomitant immunomodulators), or to the drug’s route and frequency of administration (5, 6). However, the molecular mechanisms that lead to the induction of ADA are not well understood and were initially thought to be related to the murine origin of the mAbs because they were recognized as “non-self” by the human immune system. This idea propelled the mAb discovery field to focus on engineering refined mAbs by reducing the nonhuman portions and developing chimeric, humanized, and fully human mAbs by using human libraries or humanized mice at the mAb discovery phase (7).

Unfortunately, this strategy did not abolish the immunogenicity potential of mAbs and the associated induction of ADA. The question of why and how ADA develop is further complicated by data indicating that some patients develop ADA, and some do not, and by the observation that the extent of immunogenicity may differ among patients receiving the same mAb (8). ADA that develop in patients treated with an mAb can be stratified into two main categories: 1) neutralizing ADA (*nt*ADA) that directly block and interfere with the drug’s ability to bind its target, and 2) non-neutralizing ADA (i.e., binding ADA *b*ADA) that recognize other epitopes on the drug while still retaining the mAb binding activity (9). *nt*ADA are generally considered to be more important in the clinical setting than *b*ADA because they directly reduce a drug’s efficacy. However, *b*ADA may indirectly reduce the therapeutic efficacy of an mAb by compromising bioavailability or accelerating drug clearance from the circulation. In both cases, *nt*ADA and *b*ADA substantially alter the PK and PD of the mAb being administered (10).

Originator and biosimilar tumor necrosis factor alpha (TNFα) antagonistic mAbs are used extensively in clinical settings to treat inflammatory bowel disease (IBD; e.g., Crohn’s disease and ulcerative colitis), rheumatoid arthritis, and other chronic inflammatory associated disorders such as psoriasis, psoriatic arthritis, and ankylosing spondylitis (11). TNFα antagonists help reduce inflammatory responses by targeting both membrane-bound and soluble TNFα. Neutralizing soluble TNFα prevents its binding to its receptor and impedes the secretion and upregulation of the signal cascade, thereby inhibiting its biological activity. The binding of TNFα antagonists to transmembrane TNFα on immune effector cells causes their destruction by inducing cell apoptosis or cell lysis through reverse signaling (12).

Currently, five TNFα antagonists have been approved by both the U.S. Food and Drug Administration and the European Medicines Agency: infliximab (IFX), adalimumab (ADL), etanercept, golimumab, and certolizumab pegol (2). Additionally, several biosimilars have already been approved or are in various stages of development (13). Both IFX and ADL belong to the group of TNFα antagonists and are routinely used in clinical settings to treat inflammatory diseases. IFX is a chimeric mAb (75% human and 25% murine), whereas ADL is fully human. The reported immunogenicity extent of these drugs is inconsistent. Whereas pharmaceutical companies report 10–15% and 2.6–26% immunogenicity for IFX and ADL, respectively (14), clinical data suggest higher immunogenicity rates for these drugs (15). Patients treated with IFX and ADL can be stratified based on the characteristics of their response to treatment or lack thereof. Primary non-responders are patients whose disease does not respond to the drug at all, and a certain subset of these may be mediated via early formation of ADA (15, 16). Secondary non-responders are patients who initially respond to the drug but later fail treatment, often due to development of ADA (for IFX, this was reported to develop mostly within 12 months of treatment initiation) (16).

Studies reporting immunogenicity following mAb administration and ADA prevalence have been inconsistent due in part to the various assay formats used to monitor immunogenicity in the clinic (17). Current limitations of each available format might reduce utility in clinical and research settings and complicate data interpretation. Some assays have a poor dynamic range and may generate false negative results because of interfering interaction with another circulating drug, or conversely, false positive results due to the presence of other antibodies such as rheumatoid factor (18). The pros and cons of available ADA detection assays were previously elaborated, and the formation of ADA following treatment with IFX, ADL, and other TNFα antagonists, including newly developed biosimilars, have been extensively studied and reviewed elsewhere (5, 19-21). Notwithstanding the effort invested in understanding the reasons that mAb immunogenicity and strategies to increase mAb efficacy, little is known about the molecular mechanism that governs the formation of ADA following treatment with an mAb.

In this study, we investigated the molecular landscape of ADA following treatment with TNFα antagonists. First, we developed a simple bio-immunoassay that accurately quantifies ADA levels in patient sera. We further modified the bio-immunoassay to evaluate the neutralization capacity of the ADA. Next, we aimed to profile the immune response following mAb administration. We used flow cytometry to determine the frequency of B cells in the circulation and whether the dynamics of the immune response was akin to vaccine response. Finally, we used next-generation sequencing (NGS) and high-resolution shotgun tandem mass spectrometry (LC-MS/MS) to elucidate the molecular composition of serum ADA. Using our bio-immunoassay we found that ADA levels in sera from 54 patients ranged between 2.7 and 1,268.5 μg/ml. The modified bio-immunoassay enabled us to differentiate between patients who have high and low neutralization capacity. Interestingly, we found that patients with a high neutralization capacity showed a strong bias in the λ/κ light chain ratio thereby suggesting that *nt*ADA exhibits a preference for λ light chains.

To elucidate the nature of the immune response following drug administration we chose to study a patient with IBD who was treated with IFX and who had high ADA levels and neutralization capacity. At 10 days (D10) following IFX infusion, the patient exhibited an approximately 13-fold increase in the frequency of plasmablasts (PB) and unchanged frequency of activated memory B cells (mBC), compared with the pre-infusion time point (D0). Comparative NGS analysis of the antibody heavy chain variable region (V_H_) from isolated PB at D0 and D10, showed a significant temporal decrease in the level of somatic hypermutation (SHM) and an increase in the length of the complementary determining region 3 of the antibody heavy chain (CDRH3). Moreover, the proteomic analysis of serum ADA supports the observation obtained from the neutralization capacity assays, that a preference for using λ light chains exists. These data suggest a possible mechanism whereby the humoral immune response following the administration TNFα antagonists is governed by a T cell-independent (TI) response. This response may be induced by the formation of immunocomplexes (drug-TNFα-ADA) serving as a strong driver of immunogenicity that in-turn diverts the immune response to TI pathway were B cells are activated by B cell receptor (BCR) cross-linking.

## Materials and Methods

### Over expression and purification of rhTNFα

The sequence-encoding residues Val77–Leu233 of human TNFα was cloned and fused to the N-terminal 6xHis tag in pET-28a+ vector (Novagen) and transformed into *Escherichia coli* Rosetta (DE3) cells (Novagen). A single colony was inoculated into 2ml LB supplemented with Kanamycin at final concentration of 100µg/ml and incubated over night (O.N.) at 37°C, 250 RPM. The culture was next re-inoculated into a 0.5L Erlenmeyer containing LB supplemented with Kanamycin, and grown at 37°C 250RPM until O.D._600_∼0.6-0.8 was reached. Induction was carried out by supplementing bacterial culture with IPTG (0.1mM final concentration) and incubating the culture for 3 hours at 37°C, 250RPM. Bacterial cells were harvested by centrifugation at 8000 RPM, 15 minutes, at 10°C (SORVALL RC6 Plus, Thermo Fisher Scientific) and cell pellet was stored O.N. at −20°C. Next, pellet was re-suspended in 30ml of binding buffer (50mM sodium phosphate buffer pH 8.0, 300mM NaCl, 10mM imidazole) and sonicated on ice for 8 cycles of 30 seconds pulse with 2-minute pause (W-385 sonicator, Heat Systems Ultrasonics). Following sonication, cells were centrifuged at 12000 RPM, 30 minutes, 4°C (SORVALL RC 6+) and supernatant was applied to a HisTrap affinity column (GE Healthcare) that was pre-equilibrated with binding buffer. All affinity purification steps were carried out by connecting the affinity column to a peristaltic pump with flow rate of 1/ml/min. Column was washed with 5 column volumes (CV) of wash buffer (50mM Sodium phosphate, pH 8.0, 300mM NaCl, 10% glycerol, 20mM imidazole) followed by elution of rhTNFα with 5CV of elution buffer (50mM Sodium phosphate, pH 6.0, 300mM NaCl, 10% glycerol, 500mM imidazole). Elution was collected in 1ml fractions and were analyzed by 12% SDS–PAGE. Fractions containing clean rhTNFα were merged and dialyzed using Amicon Ultra (Mercury) cutoff 3K against PBS (pH 7.4). Dialysis products were analyzed by 12% SDS–PAGE for purity and concentration was measured using Take-5 (BioTek Instruments). To test functionality of the produced rhTNFα, 96 well plate (Nunc MaxiSorp™ flat-bottom, Thermo Fisher Scientific) was coated with 1μg/ml (in PBS) of purified rhTNFα and commercial hTNFα (PHC3011, Thermo Fisher Scientific) and incubated at 4°C O.N. ELISA plates were then washed three times with PBST (0.1% v/v Tween 20 in PBS) and blocked with 300µl of 2% w/v BSA in PBS for 1 hour at 37°C. Next, ELISA plates were washed three time with PBST, and incubated for 1 hour, room temperature (RT) in triplicates with anti-TNFα mAb (Infliximab or Adalimumab) in 2% w/v BSA, PBS at the starting concentration of 50nM with 3-fold dilution series. Plates were then washed three times with PBST with 30 second incubation time at each washing cycle. For detection, 50μl of anti-human H+L HRP conjugated antibody (Jackson) was added to each well (1:5000 ratio in 2% w/v BSA in PBS) and incubated for 1 hour at RT, followed by three washing cycles with PBST. Developing was carried out by adding 50µl of 3,3’,5,5’-Tetramethylbenzidine (TMB, Southern Biotech) and reaction was quenched by adding 50µl 0.1M sulfuric acid. Plates were read using the Epoch Microplate Spectrophotometer ELISA plate reader (BioTek Instruments).

### Over expression and purification of IdeS

The coding sequence corresponding to amino acid residues 38–339 of *S. pyogenes* IdeS (numbered from the start of the signal sequence) was sub-cloned into the expression vector pET28b (Novagen). The coding sequencing was sub-cloned at the 3’ end of Thioredoxin 6xHis-TEV. The complete construct was sub-cloned as previously described (22) and was kindly donated by Dr. Ulrich von Pawel-Rammingen from the Department of Molecular Biology, Umea University. The transformation of pET-TRX_b plasmid harboring the IdeS encoding gene (pET-IdeS) was carried out as follows: 200μl of chemical-competent *E. coli* BL21-DE3 cells were thawed on ice for 20 minutes. 50ng of the plasmid pET-IdeS was added to the thawed competent cells and incubated on ice for 20 minutes with gentle mixing every 5 minutes. Next, heat shock was applied by incubating the cells at 42°C for 2 minutes followed by incubation on ice water for 2 minutes. For phenotypic expression, 800μl of LB was added, and cells were incubated at 37°C, 250 RPM for 1 hour in a horizontal position. Cells were plated on LB agar supplemented with Kanamycin and incubated at 37°C overnight (O.N). Single colony was inoculated into 2ml LB supplemented with Kanamycin and incubated O.N. at 37°C, 250 RPM. Next day, 2ml from the grown cultures were inoculated into two 2liter flasks, each containing each 500ml LB supplemented with Kanamycin. Over expression and purification of IdeS was carried out as described for rhTNFα with a minor modification as follow: Ides was eluted with imidazole gradient (50, 150, 500mM imidazole), total of 20ml. 20 fractions of 1ml were collected from each elution step and evaluated for their purity using 12% SDS–PAGE. All fractions containing clean IdeS were merged and dialyzed O.N. at 4°C against 4L of PBS (pH 7.4), using SnakeSkin dialysis tubing with 10 kDa cutoff (Thermo Fisher Scientific). Dialysis products were analyzed by 12% SDS–PAGE.

### Production of mAb-F(ab’)_2_

Intact clinical grade IFX or ADL (designated here as mAb) were digested using in-house produced IdeS. 10mg of mAb was incubated with 300µg of IdeS in the final volume of 500µl PBS for 2.5 hours at 37°C, followed by a spike-in of additional 300µg of IdeS to achieve full digestion of the Fc fragments. IdeS inactivation was carried out by adding 0.1M of citric acid pH 3 and incubation for 1 minute at RT followed by the addition of PBS (pH 7.4) to neutralized acidic pH. Next, reaction mixture was applied to a 1 mL HiTrap KappaSelect affinity column (GE Healthcare Life Sciences). All affinity purification steps were carried out by connecting the affinity column to a peristaltic pump with flow rate of 1ml/min. The reaction mixture was recycled three times through the KappaSelect column to maximize the capture of intact mAb and mAb-F(ab’)_2_. KappaSelect column was subsequently washed with 5CV of PBS and eluted with 10CV of 100mM glycine·HCl (pH 2.7). Collected 1ml elution fractions were immediately neutralized with 100µl of 1.5M Tris·HCl (pH 8.8). Next, the recovered intact mAb and mAb-F(ab’)_2_ fragments were applied to a custom packed 1ml Protein-G agarose column (GenScript). The reaction mixture was recycled three times through the column, which was subsequently washed with 5CV of PBS and eluted with 10CV of 100mM glycine·HCl (pH 2.7). The 10ml elution fraction was immediately neutralized with 1ml of 1.5M Tris·HCl (pH 8.8). The recovered 10ml mAb-F(ab’)_2_ fragments were dialyzed overnight at 4°C against 4L of PBS (pH 7.4) using SnakeSkin dialysis tubing with 10kDa cutoff (Thermo Fisher Scientific). Recovered mAb-F(ab’)_2_ sample were evaluated for purity by SDS-PAGE and their concentration measured by Take5 (BioTek instruments).

To test the functionality of the produced mAb-F(ab’)_2_, 96 ELISA plates (Nunc MaxiSorp™ flat-bottom, Thermo Fisher Scientific) were coated with 1µg/ml of rhTNFα in PBS and incubated at 4°C O.N. ELISA plates were then washed three times with PBST and blocked with 300µl of 2% w/v BSA in PBS for 1 hour at 37°C. Next, 50nM of intact mAb and mAb-F(ab’)_2_ (IFX or ADL) in blocking solution was added to each well in triplicates in a 3 fold dilution series, and plates were incubated at RT for 1 hour. Next, plates were washed three times with PBST with 30 second incubation time at each washing cycle. For detection, HRP conjugated anti-human kappa light chain (Jackson) was added to each well (50μl, 1:5000 ratio in 2% w/v BSA in PBS) and incubated for 1 hour at RT, followed by three washing cycles with PBST. Developing was carried out by adding 50µl of TMB and reaction was quenched by adding 0.1M sulfuric acid. Plates were read using the Epoch Microplate Spectrophotometer ELISA plate reader. To evaluate the purity of the mAb-Fa(b’)_2_ samples (i.e. to make sure there are no traces of intact antibody or Fc fragment in the sample), 96 ELISA plate (Nunc MaxiSorp™ flat-bottom, Thermo Fisher Scientific) were coated with 5µg/ml of intact mAb and mAb-F(ab’)_2_ in PBS and incubated at 4°C O.N. Next, plates were washed three time with PBST and blocked with 300µl 2% w/v BSA in PBS for 1 hour at 37°C. Next, plates were washed three times with 300 µl PBST, followed by the incubation with HRP conjugated anti-human IgG Fc antibody (Jackson) diluted 1:5000 in PBST. Developing was carried out by adding 50µl of TMB and reaction was quenched by adding 0.1M sulfuric acid. Plates were read using the Epoch Microplate Spectrophotometer ELISA plate reader (BioTek Instruments).

### Generation of ADA standard

A pool of 17 ADA to IFX positive sera were collected at Sheba Medical Center, and passed through a 2ml custom packed protein G agarose column (GenScript). The pooled sera was recycled three times over the column, which was subsequently washed with 5CV of PBS and eluted with 10CV of 100mM glycine·HCl (pH 2.7). The 10ml elution fraction was immediately neutralized with 1ml of 1.5M Tris·HCl (pH 8.8). The purified mAbs were immediately passed over a custom made rhTNFα affinity column (NHS-activated agarose beads, Thermo Fisher Scientific) in gravity mode. The purified mAbs were recycled three times over the column, which was subsequently washed with 5CV of PBS and eluted with 10CV of 100mM glycine·HCl (pH 2.7). The 10ml elution fraction was immediately neutralized with 1ml of 1.5M Tris·HCl (pH 8.8). The purified mAbs were dialyzed overnight at 4°C against 4L of PBS (pH 7.4) using SnakeSkin dialysis tubing with 10kDa cutoff (Thermo Fisher Scientific). Purified mAbs were analyzed for purity using 12% SDS-PAGE and concentration was determined by Take3 (BioTek instruments).

To test functionality, 96 ELISA plate were coated with 5µg/ml of mAb-F(ab’)_2_ in PBS (pH 7.4) and incubated at 4°C O.N. ELISA plates were then washed three times with PBST and blocked with 300µl of 2% w/v BSA in PBS for 1 hour at 37°C. Next, 50nM of the purified ADA in blocking solution were added to each well in triplicates with 3-fold dilution series and plates were incubated at RT for 1 hour. Next, plates were washed three times with PBST with 30 second incubation time at each washing cycle. Next, anti-human Fc HRP conjugate (Jackson) was added to each well at the detection phase (50μl, 1:5000 ratio in 2% w/v BSA in PBS) and incubated for 1 hour at RT, followed by three washing cycles with PBST. Developing was carried out by adding 50µl of TMB and reaction was quenched by adding 0.1M sulfuric acid. Plates were read using the Epoch Microplate Spectrophotometer ELISA plate reader.

### Quantitative measurement of ADA in serum

The schematic configuration of the bio-immunoassay for the quantitative measurement of ADA in serum is described in Fig. 3B and was carried out as follows: ELISA plates that were coated overnight at 4°C with 5μg/ml produced IFX-F(ab’)_2_ in PBS (pH 7.4). ELISA plates were then washed three times with PBST and blocked with 300µl of 2% w/v BSA in PBS for 1 hour at 37°C. Next, triplicates of 1:400 diluted serum samples were added at triplicates and serially diluted 2 fold in 2% w/v BSA in PBS, 10% horse serum (Biological Industries) and 1% Tween 20 in PBS (1:400– 1:51,200 serum dilution factor). Plates were incubated for 1 hour at RT. On the same plate, serial dilutions of 10nM ADA standard were incubated in triplicate and serially diluted 2 fold in 2% w/v BSA in PBS, 10% horse serum (Biological Industries) and 1% Tween 20 in PBS, to allow the conversion of the tested serum to units per milliliter. ELISA plates were washed three times with PBST and 50μl of HRP conjugated anti-human IgG Fc was added to each well (50μl, 1:5000 ratio in 2% w/v BSA in PBS) and incubated for 1 hour at RT. ELISA plate was then washed three times with PBST and developed by adding 50µl of TMB followed by quenching with 50μl 0.1M sulfuric acid. Plates were read using the Epoch Microplate Spectrophotometer ELISA plate reader.

**Figure 1:**
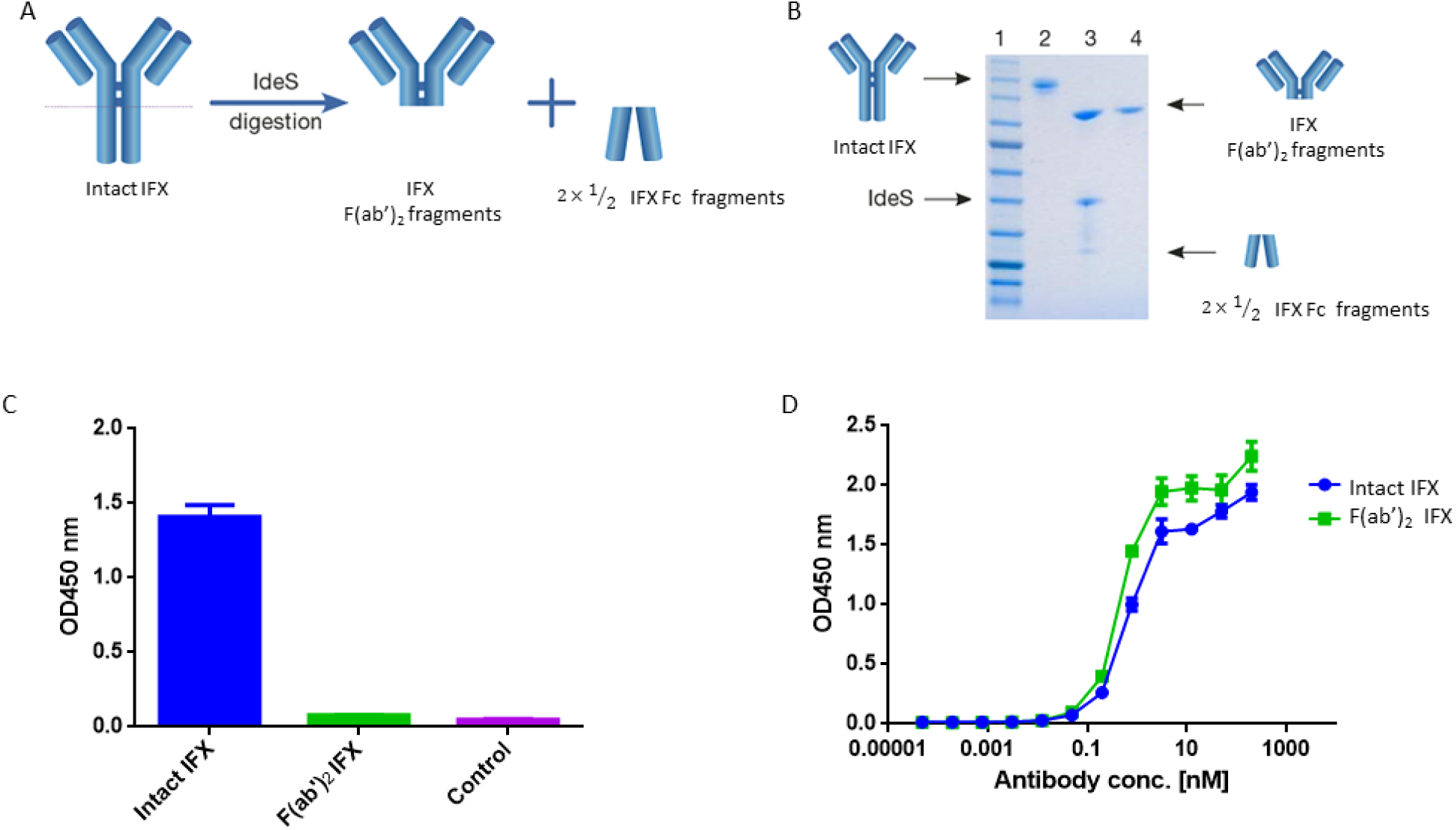
IFX digestion and IFX-F(ab’)_2_ purification. (A) Schematic representation of IgG digestion with IdeS. IdeS is a highly specific immunoglobulin-degrading enzyme that cleaves below the disulfide bonds in the IgG hinge region. The cleavage results in the production of IFX-F(ab’)_2_ fragment and two 1/2 Fc fragments. (B) SDS-PAGE analysis of intact IgG (lane 2), following IdeS digestion (lane 3) and purified IFX-F(ab’)_2_ following a 2-step affinity chromatography purification including protein A and kappa-select columns (lane 4). (C) Presence of Fc and intact IgG traces was measured by direct ELISA where intact IFX and purified IFX-F(ab’)_2_ were compared to a control antigen (streptavidin) as coating agents followed by direct incubation with an anti-Fc HRP conjugate at the detection phase. (D) The functionality of the recovered IFX-F(ab’)_2_ was confirmed by testing it for TNFα binding by ELISA in comparison to intact IFX. The ELISA setup included TNFα as the coating agent and anti-κ HRP conjugate at the detection phase. For panel C–D, triplicate averages were calculated as mean, with error bars indicating s.d.

**Figure 2:**
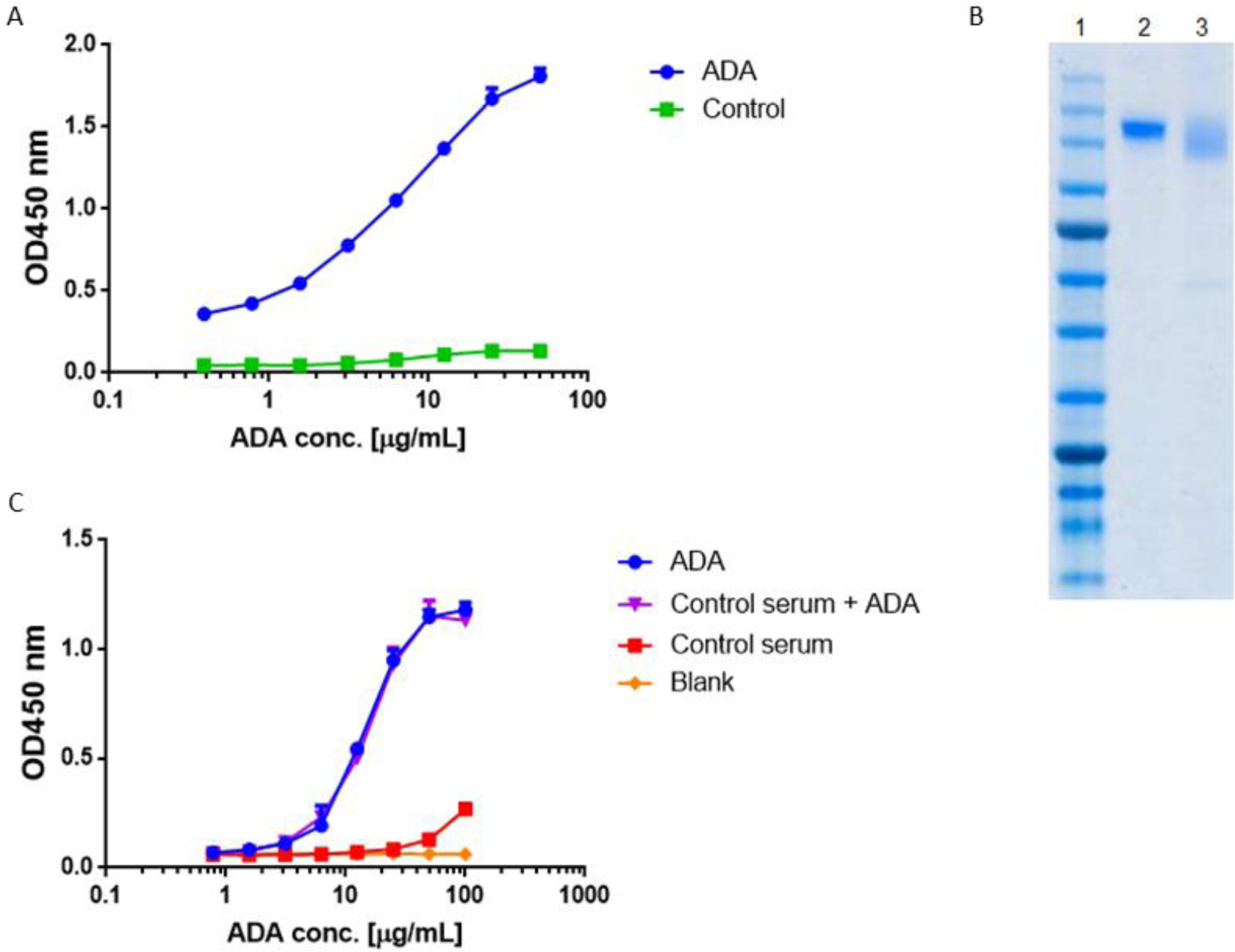
Standard curve for ADA quantification in patients treated with IFX. ADA were purified from sera of 17 patients treated with IFX, utilizing consecutive affinity chromatography steps including protein G and custom made IFX-F(ab’)_2_ columns. (A) Purified ADA were tested in ELISA for functionality. TNFα was used as the coating agent followed by incubation with purified ADA and anti-Fc HRP conjugate at the detection phase. Control included serum obtained from a healthy donor. (B) SDS-PAGE analysis of intact IFX (lane 2) and purified ADA (lane 3). (C) The effect of serum on ADA standard was tested in ELISA by s piking-in differential concentrations of ADA into ADA negative serum.

**Figure 3:**
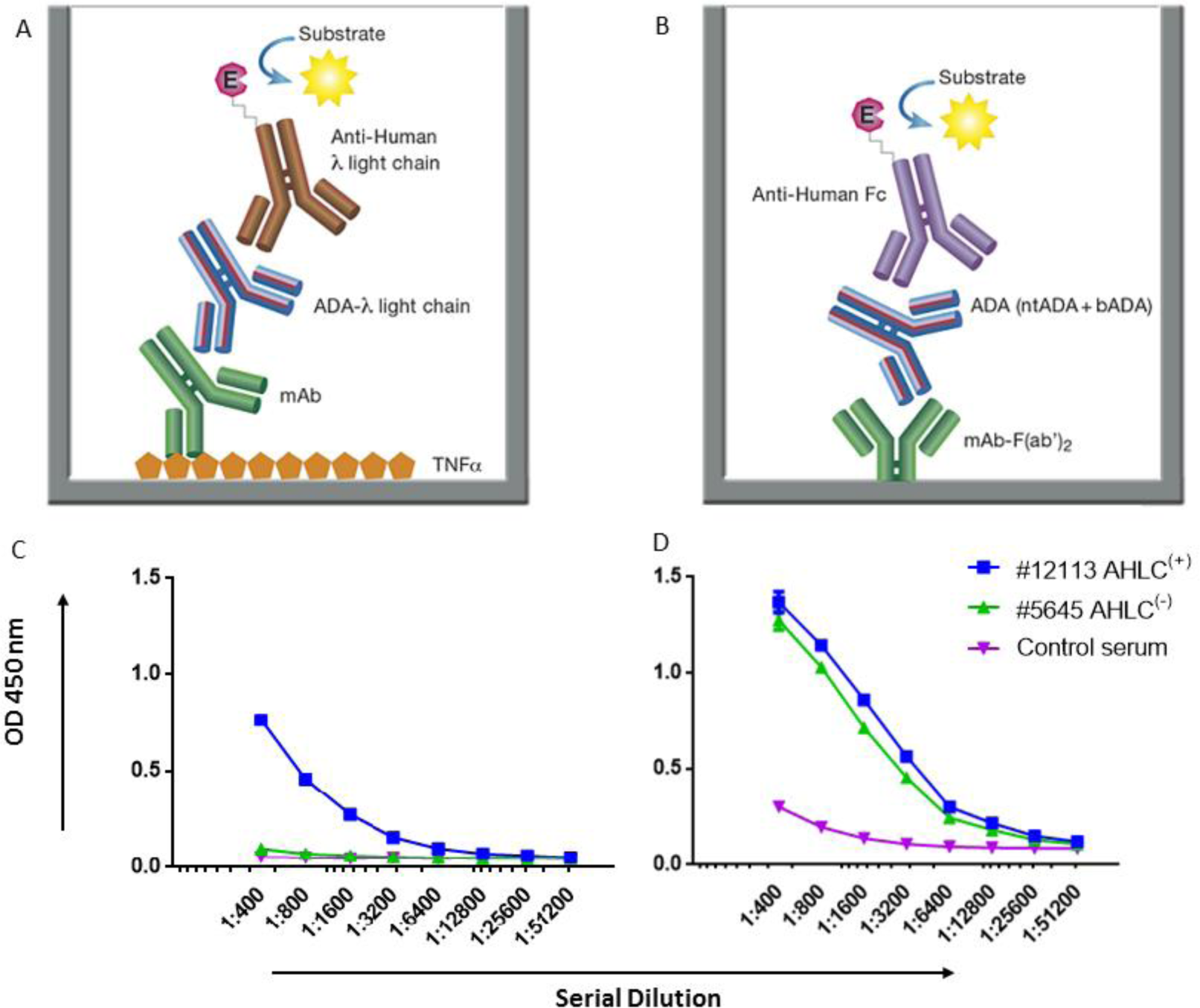
AHLC and the newly developed mAb-F(ab’)_2_ based bio-immunoassay configuration and their application on serum samples from patients treated with IFX. (A) AHLC assay is based on an ELISA where TNFα is used as the coating agent, following the incubation with the mAb drug followed by serial dilutions of the tested sera. anti-λ HRP conjugate is used at the detection phase. (B) The newly developed mAb-F(ab’)_2_ based bio-immunoassay configuration. The assay is based on an ELISA where mAb-F(ab’)_2_ is used as the coating agent followed by serial dilutions of the tested sera. Anti-Fc HRP conjugate is used at the detection phase. (C) ELISA obtained by utilizing the AHLC assay on two serum samples. Using this assay, one of the tested sera showed detectable levels of ADA (AHLC^(+)^) and one had no detectable levels of ADA (AHLC^(-)^). (D) Both serum samples were tested by the newly developed mAb-F(ab’)_2_ based bio-immunoassay. This assay was able to detect ADA in both sera. For C–D, averages were calculated as mean from triplicates, with error bars indicating s.d.

### Neutralization index of ADA

Neutralization capacity was determined using ELISA plates that were coated overnight at 4°C with 5μg/ml IFX-F(ab’)_2_ in PBS (pH 7.4). Next, coating solution was discarded and ELISA plates were blocked with 300µl of 2% w/v BSA in PBS for 1 hour at 37°C. Blocking solution was discarded and 50µl of 200 nM rhTNFα in 2% w/v BSA were added to the positive rhTNFα wells, and 2% w/v BSA in PBS was added to the negative rhTNFα wells for 1 hour at RT. Next, triplicates of 1:400 diluted serum samples with/without 200nM rhTNFα were added to the positive/negative rhTNFα wells (respectively) and serially diluted 2-fold in 2% w/v BSA, 10% horse serum (Biological Industries) and 1% Tween 20 in PBS (1:400– 1:51,200 serum dilution factor). Plates were incubated for 1 hour at RT. ELISA plates were washed three times with PBST and 50μl of HRP conjugated anti-human IgG Fc antibody or anti HRP conjugated His-tag antibody were added at the detection phase (50μl, 1:5000 ratio in 2% w/v BSA in PBS) and incubated for 1 hour at RT, followed by three washing cycles with PBST. Developing was carried out by adding 50µl of TMB and reaction was quenched by adding 0.1M sulfuric acid. Plates were read using the Epoch Microplate Spectrophotometer ELISA plate reader.

Neutralization index was calculated as following: an ELISA equation curve was calculated separately for wells with and without rhTNFα, using the GraphPad Prism software. The average triplicate signal which are 3 × standard deviation above the background signal was substituted in the ELISA equation curve to extract the serial dilution value. The logarithmic difference of the value with/without rhTNFα represents the neutralization index.

### Blood processing

IFX treated patients with IBD cared for in the Department of Gastroenterology at the Sheba medical center were included in the study. All subjects signed an informed consent, and the study was approved by the Ethics Committee of the medical center. All patients received IFX on a scheduled regimen and blood samples were drawn immediately before their scheduled IFX infusion. Blood was collected into a single Vacutainer Lithium Heparin collection tube (BD Bioscience).

For NGS analysis, blood was collected from a male donor treated with IFX, before IFX administration and 10 days after administration. 30ml of peripheral blood were collected into 3 single Vacutainer K-EDTA collection tubes (BD Biosciences). Collection of peripheral blood mono-nuclear cells (PBMCs) was performed by density gradient centrifugation, using Uni-SepMAXI+ lymphocyte separation tubes (Novamed) according to the manufacturer’s protocol.

### Fluorescence-Activated Cell Sorting Analysis and sorting of B cell populations

PBMCs were stained for 15 minutes in cell staining buffer (BioLegend) at RT in the dark using the following antibodies: anti-CD3–PerCP (clone OKT3; BioLegend), anti-CD19– Brilliant Violet 510 (clone HIB19; BioLegend), anti-CD27–APC (clone O323; BioLegend), anti-CD38– APC-Cy7 (clone HB-7; BioLegend), and anti-CD20–FITC (clone 2H7; BioLegend).

The following B cell population was sorted using a FACSAria cell sorter (BD Bioscience): CD3−CD19+CD20-CD27++CD38^high^

B cell subpopulations were sorted and collected into TRI Reagent solution (Sigma Aldrich) and frozen at −80°C.

### Amplification of V_H_ and V_L_ repertoires from B cells

Total RNA was isolated using RNeasy micro Kit (Qiagen), according to manufacturer’s protocol. First-strand cDNA generation was performed with 100ng of isolated total RNA using a SuperScript RT II kit (Invitrogen) and oligo-dT primer, according to manufacturer’s protocol. After cDNA synthesis, PCR amplification was performed to amplify the V_H_ and V_L_ genes using a primer set described previously (23) with overhang nucleotides to facilitate Illumina adaptor addition during the second PCR (Table S1). PCR reactions were carried out using FastStart™ High Fidelity PCR System (Roche) with the following cycling conditions: 95°C denaturation for 3 min; 95°C for 30 sec, 50°C for 30 sec, and 68°C for 1 min for four cycles; 95°C for 30 sec, 55°C for 30 sec, and 68°C for 1 min for four cycles; 95°C for 30 sec, 63°C for 30 sec, and 68°C for 1 min for 20 cycles; and a final extension at 68°C for 7 min. PCR products were purified using AMPure XP beads (Beckman Coulter), according to manufacturer’s protocol (ratio × 1.8 in favor of the beads). Recovered DNA products from the first PCR was applied to a second PCR amplification to attach Illumina adaptors to the amplified V_H_ and V_L_ genes using the primer extension method as described previously (24). PCR reactions were carried out using FastStart™ High Fidelity PCR System (Roche) with the following cycling conditions: 95°C denaturation for 3 min; 95°C for 30 sec, 40°C for 30 sec, and 68°C for 1 min for two cycles; 95°C for 30 sec, 65°C for 30 sec, and 68°C for 1 min for 7 cycles; and a final extension at 68°C for 7 min. PCR products were applied to 1% agarose DNA gel electrophoresis and gel-purified with Zymoclean™ Gel DNA Recovery Kit (Zymo Research) according to the manufacturer’s instructions. V_H_ and V_L_ libraries concentration were measured using Qubit system (Thermo Fisher Scientific) and library quality was assessed using the Bioanalyzer 2100 system (Agilent) or the 4200 TapeStation system (Agilent). All V_H_ libraries were produced in duplicates starting with RNA as the common source template. The V_L_ were produced with one replicate.

V_H_ and V_L_ libraries from sorted B cell were subjected to NGS on the MiSeq platform with the reagent kit V3 2×300 bp paired-end (Illumina), using an input concentration of 16pM with 5% PhiX.

Raw fastq files were processed using our recently reported ASAP webserver (25). ASAP analysis resulted in a unique, full-length V_H_ and V_L_ gene sequences database for each time point. The resultant database was used as a reference database to search the LC-MS/MS spectra.

### Proteomic Analysis of the Serum ADA to IFX

Total IgG from each time point (D0, D10) were purified from 7-10ml of serum by protein G enrichment. Serum was diluted 2 fold and passed through a 5ml Protein G agarose column (GeneScript). The diluted serum was recycled three times over the column, which was subsequently washed with 10CV of PBS and eluted with 7CV of 100mM glycine·HCl (pH 2.7). A total of 35 fractions of 1ml were collected and immediately neutralized with 100µl of 1.5M Tris·HCl (pH 8.8). All elution fractions were evaluated for their purity using 12% SDS–PAGE and 11 purified 1 ml IgG fractions were combined and dialyzed overnight at 4°C against 4L of PBS (pH 7.4) using SnakeSkin dialysis tubing with 1 kDa cutoff (Thermo Fisher Scientific).

Next, 9mg of total IgG were digested with 100µg of IdeS in the final volume of 2ml PBS for 5 hour at 37°C. IdeS inactivation was carried out by adding 0.1M of citric acid pH 3 and incubation for 1 minute at RT followed by the addition of PBS (pH 7.4) to neutralize the low pH. Total serum F(ab’)_2_ was then applied to a one ml custom made affinity column comprised of IFX-F(ab’)_2_ coupled to NHS-activated agarose beads (Thermo Fisher Scientific). The purified serum F(ab’)_2_ were recycled three times over the affinity column, which was subsequently washed with 5CV of PBS and eluted with 15CV of 100mM glycine·HCl (pH 2.7) and collected into Maxymum Recovery Eppendorf (Axygen Scientific). A total of 30×0.5ml elution fractions and 1×50ml flow-through were immediately neutralized with 50 and 100µl (respectively) of 1.5M Tris·HCl (pH 8.8). The purified antigen-specific F(ab’)_2_ were dialyzed overnight at 4°C against 4L of PBS (pH 7.4) using SnakeSkin dialysis tubing with 10kDa cutoff (Thermo Fisher Scientific). Elution and flow-through fractions were trypsin-digested, and resulting peptides were fractionated and sequenced by nanoflow LC-electrospray MS/MS on an Orbitrap Velos Pro hybrid mass spectrometer (Thermo Scientific), in the UT Austin mass spectrometry core facility as described previously (26). MS/MS raw files were analyzed by MaxQuant software version 1.6.0.16 (27) using the MaxLFQ algorithm (28) and peptide lists were searched against the common contaminants database by the Andromeda search engine (29) and a custom protein sequence database consisting of the donor-specific V_H_ and V_L_ sequences derived from NGS of individual donor B cells. All searches were carried out with cysteine carbamidomethylation as a fixed modification and methionine oxidations as variable modifications. The false discovery rate was set to 0.01 for peptides with a minimum length of seven amino acids and was determined by searching a reverse decoy database. Enzyme specificity was set as C-terminal to arginine and lysine as expected using trypsin as protease, and a maximum of two missed cleavages were allowed in the database search. Peptide identification was performed with an allowed initial precursor mass deviation up to 7ppm and an allowed fragment mass deviation of 20ppm. For LFQ quantification the minimal ratio count was set to 2, and match between runs was performed with three mass-spec injections originating from the same sample. MaxQunat output analysis file, “peptides.txt”, was used for further processing. Total peptides that were identified in the elution samples were filtered using the following criteria: (a) were not identified as contaminates; (b) did not match to the reversed decoy database; (c) were identified as peptides derived from the region comprising the CDRH3, J region, FR4 and the ASTK motif (derived from the N-terminal of the C_H_1 region). The CDRH3 derived peptides were further characterized as informative CDRH3 peptides (*i*CDRH3 peptides) only if they map exclusively to a single antibody clonotype. A clonotype was defined as all sequences that comprise CDRH3 with the same length and identity tolerating one amino acid mismatch, and same V, J family. The intensities of high confidence *i*CDRH3 peptides were averaged between replicates while including only peptides that were observed in at least two out of the three replicates. Clonotype frequencies within each sample were calculated using only *i*CDRH3 peptides and were determined to be antigen-specific if their frequency in the elution fraction was at least 5 fold greater than their frequency in the flow-through fraction. The CDRH3 sequences identified by the mapping of high confidence MS/MS peptides were used to generate a complete list of full length VH sequences. These VH sequences were used to analyze the repertoire measures of the antibodies that were identified in the donors’ serum.

Same filtering criteria was applied to peptides derived from the constant region of both κ and λ light chains. By quantifying the accumulative intensities of these peptides, we calculated the ratio of κ:λ light chain from antibodies that were derived from the affinity column elution fraction which represent both *nt*ADA and *b*ADA, and in the affinity column flow through fraction which represent the depleted ADA fraction.

### Study population

IFX and ADL treated patients with IBD cared for in the Departments of Gastroenterology at Sheba medical center were included in the study. All subjects signed an informed consent, and the study was approved by the Ethics Committee of Sheba medical center. IFX and ADL and ADA serum levels were routinely measured at trough immediately before infusion. All patients received IFX and ADL on a scheduled regimen. All patients that were included in this study exhibited low through levels of IFX and ADL.

### Statistical analysis

All curves were fitted on a sigmoidal dose–response curve and EC50 of each was calculated. Mann-Whitney test was used to compare continuous variables. All reported P values were two-tailed, and a P value less than 0.05 were considered statistically significant. All statistics were performed with GraphPad Prism software (version 7, San Diego, California).

## Results

### Production of mAb-F(ab’)_2_ to be used in the bio-immunoassay

To investigate the molecular landscape of ADA following mAb administration we first aimed to develop an accurate, sensitive, robust bio-immunoassay to determine ADA levels in sera. The working hypothesis was that anti-idiotypic antibodies dominate the ADA compartment (21) thus, the developed bio-immunoassay was based on the drugs’ F(ab’)_2_ portion to be used as the antigen (i.e. coating agent).

To achieve this, we used the immunoglobulin G (IgG)-cleaving enzyme (IdeS), a cysteine proteinase enzyme that proteolytically cleaves immunoglobulins below the hinge region (30) (Figure 1A). IFX was digested using IdeS by incubating 01 mg of clinical grade mAb with IdeS to reach near complete digestion. Next, IFX-F(ab’)_2_ was purified from Fc regions and undigested full IFX by consecutive affinity chromatography steps comprising protein A and kappaSelect columns.

Recovered IFX-F(ab’)_2_ purity was evaluated by SDS-PAGE (Figure 1B) and ELISA (Figure 1C) to ensure that the IFX-F(ab’)_2_ exhibits no traces of IFX-Fc/undigested IFX that will contribute to the background level when using anti-Fc HRP conjugate at the detection phase. Recovered IFX-F(ab’)_2_ samples were found to be highly pure with basal anti-Fc signal levels similar to the signal observed in the control samples. The produced IFX-F(ab’)_2_ was tested for functionality by measuring its TNFα binding capacity, using ELISA with TNFα as the coating agent, and was found to show similar functionality as that of the intact IFX (Figure 1D). ADL was subjected to the same preparative pipeline and demonstrated similar results (Figure S1).

### ADA Standard curve

Quantification of total ADA in serum requires a standard reference. Thus, we generated a standard ADA pool that facilitates the quantification of ADA levels in sera of patient treated with IFX. ADA were pooled from several serum samples collected from patients treated with IFX and purified by consecutive affinity chromatography steps comprising protein G and a custom-made IFX-F(ab’)_2_ affinity columns. We confirmed the affinity enrichment of ADA by applying the affinity chromatography elution fraction to ELISA with IFX-F(ab’)_2_ as the coated antigen (Figure 2A). The purity and concentration the recovered ADA were determined by SDS-PAGE (Figure 2B) and nanodrop.

Maximal serum concentration used in a bio-immunoassay (e.g. serum diluted 1:100 or 1:200) is a major factor that may contribute to high background signal levels due to non-specific binding. Screening several maximal serum dilutions showed that 1:400 initial serum dilution demonstrates the lowest background signal (data not shown). To evaluate if serum will affect the signal obtained from purified ADA, we spiked-in purified ADA into negative control serum that was diluted 1:400 in PBS. Serial dilution of spiked-in ADA and purified ADA showed similar signal in ELISA (Figure 2C) indicating that serum does not bias the ADA detection in our developed bio-immunoassay.

### Quantitative measurement of ADA in serum

ADA detection is technically challenging as both the analyte and antigen are antibodies which may result in the inability to differentiate between the mAb and ADA. To overcome this challenge, many assays were previously developed (5). One of these immunoassays is the anti-human λ chain (AHLC) immunoassay that is used in clinical setups for monitoring the formation of ADA (31). The principle of this assay is to detect ADA comprising λ light chain, thus avoiding cross reactivity with the drug that comprises the κ light chain (Figure 3A).

While AHLC is suitable for monitoring the development of ADA in clinical setups, when one aims to study the molecular composition of ADA there is a need to provide quantitative measures of total ADA in serum. Thus, we developed a new bio-immunoassay based on the F(ab’)_2_ portion of the mAb. The bio-immunoassay setup is described in Figure 3B and is based on mAb-F(ab’)_2_ as the coating antigen and anti-Fc HRP conjugate used as the detection antibody. Each of the experimental setups to test ADA in serum included serum from a healthy donor as a control and ADA standard for the quantitation of total ADA.

First, we applied the newly developed bio-immunoassay on two serum sample groups: one negative and one positive for ADA as determined by the AHLC assay (AHLC^(-)^ and AHLC^(+)^, respectively). We also included serum from a healthy subject to serve as a control for the assay specificity (i.e. serum from a subject that was not exposed to IFX). As shown in Figure 3C-D, the ELISA signals obtained when utilizing the new bio-immunoassay were higher compared to the signal obtained with the AHLC assay. Moreover, applying the new bio-immunoassay on the AHLC^(-)^ serum (no detected ADA with the AHLC assay) detected relatively high levels of ADA. These results indicate that not all ADA were detected with the AHLC assay as this assay is based on the detection of ADA that comprise the λ light chain only.

Next, to extend and generalize the above results, sera from 54 patients treated with IFX were collected at the Chaim Sheba Medical Center and tested for drug levels and ADA using the AHLC assay. The established cohort showed very low drug trough levels and based on the AHLC results, sera were stratified into two groups: 25 serum samples were identified as AHLC^(-)^ and 29 as AHLC^(+)^. Using our newly developed quantitative bio-immunoassay, we found that ADA levels in tested sera ranged between 1.82 to 1268.5 μg/ml. Serum ADA levels using AHLC compared to the new bio-immunoassay are summarized in Table 1. More importantly, the new bio-immunoassay demonstrated improved sensitivity compared to AHLC assay manifested in the detection of higher concentrations of ADA in 46 out of the 54 serum samples, of which 17 out of the 54 samples, belong to the AHLC^(-)^ group. Overall, the average fold increase in ADA detection using the new bio-immunoassay compared to the AHLC assay was 14.13 and 53.26 for the AHLC^(+)^ and AHLC^(-)^ groups, respectively.

**Table 1:**
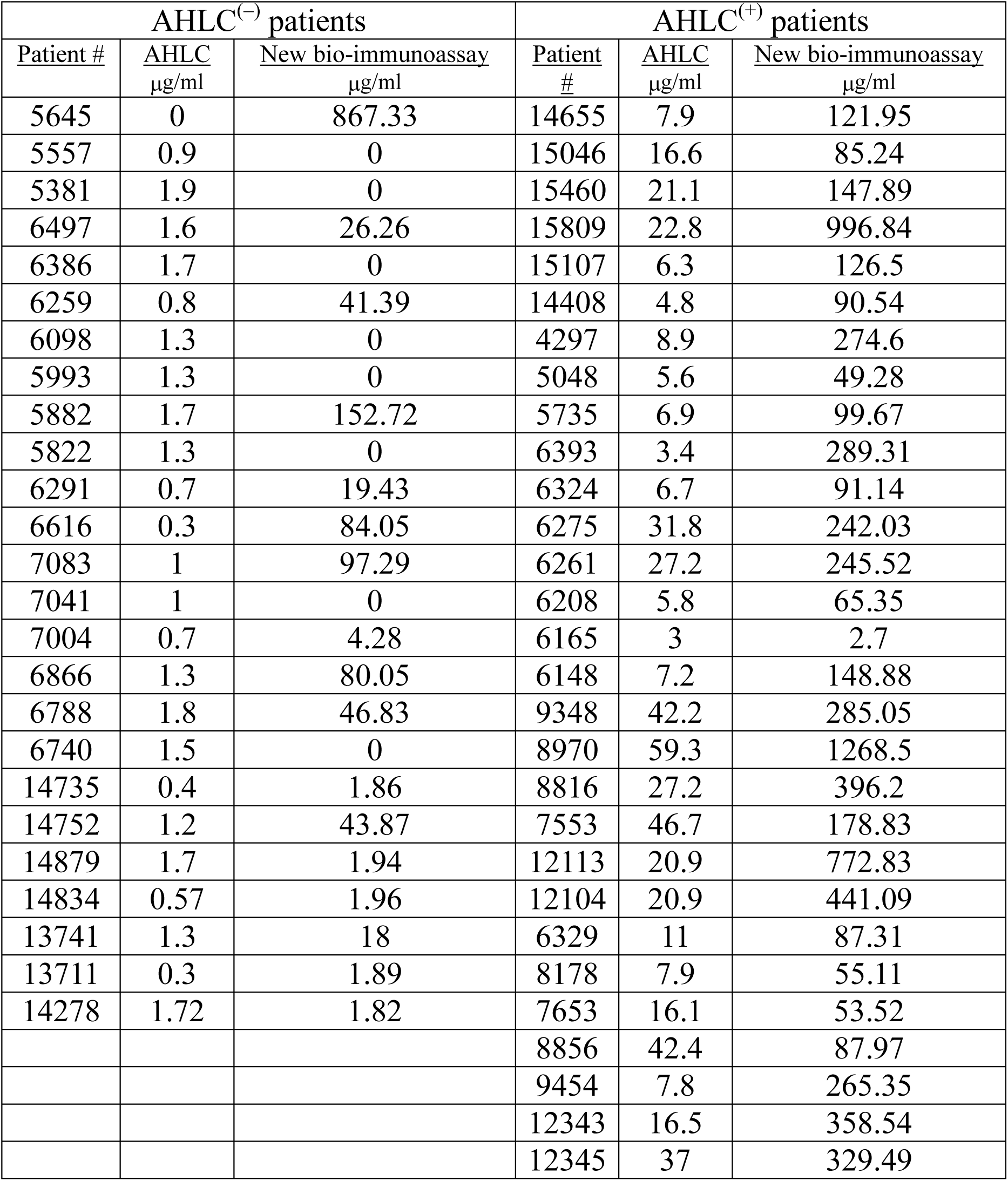
ADA concentrations in 55 serum samples from patients treated with IFX. Serum samples were initially stratified into AHLC (+) and AHLC (-) based on the AHLC assay used in the clinic. The newly developed bio-immunoassay for the quantification of total ADA was applied on all serum samples and concentration are listed. All ADA concentrations are in µg/ml.

### Neutralization index of ADA

Due to high clinical relevance and different mechanism of action of *b*ADA and *nt*ADA, identifying their relative abundances in serum can provide valuable insights regarding the nature of the immune response following mAb administration. We therefore modified our newly developed mAb-F(ab’)_2_ based bio-immunoassay by blocking the coated IFX-F(ab’)_2_ binding site with TNFα in order to obtain a differential signal compared to the unblocked assay (Figure 4A). In order to block the binding site of IFX-F(ab’)_2_ towards TNFα and prevent the binding of anti-idiotypic ADA (i.e. *nt*ADA) to the drug, recombinant human TNFα (rhTNFα) fused to a His-tag was cloned and expressed (see materials and methods). In-house production of rhTNFα was essential, as the N terminal His-tag was used for monitoring the presence of the rhTNFα throughout the bio-immunoassay.

**Figure 4:**
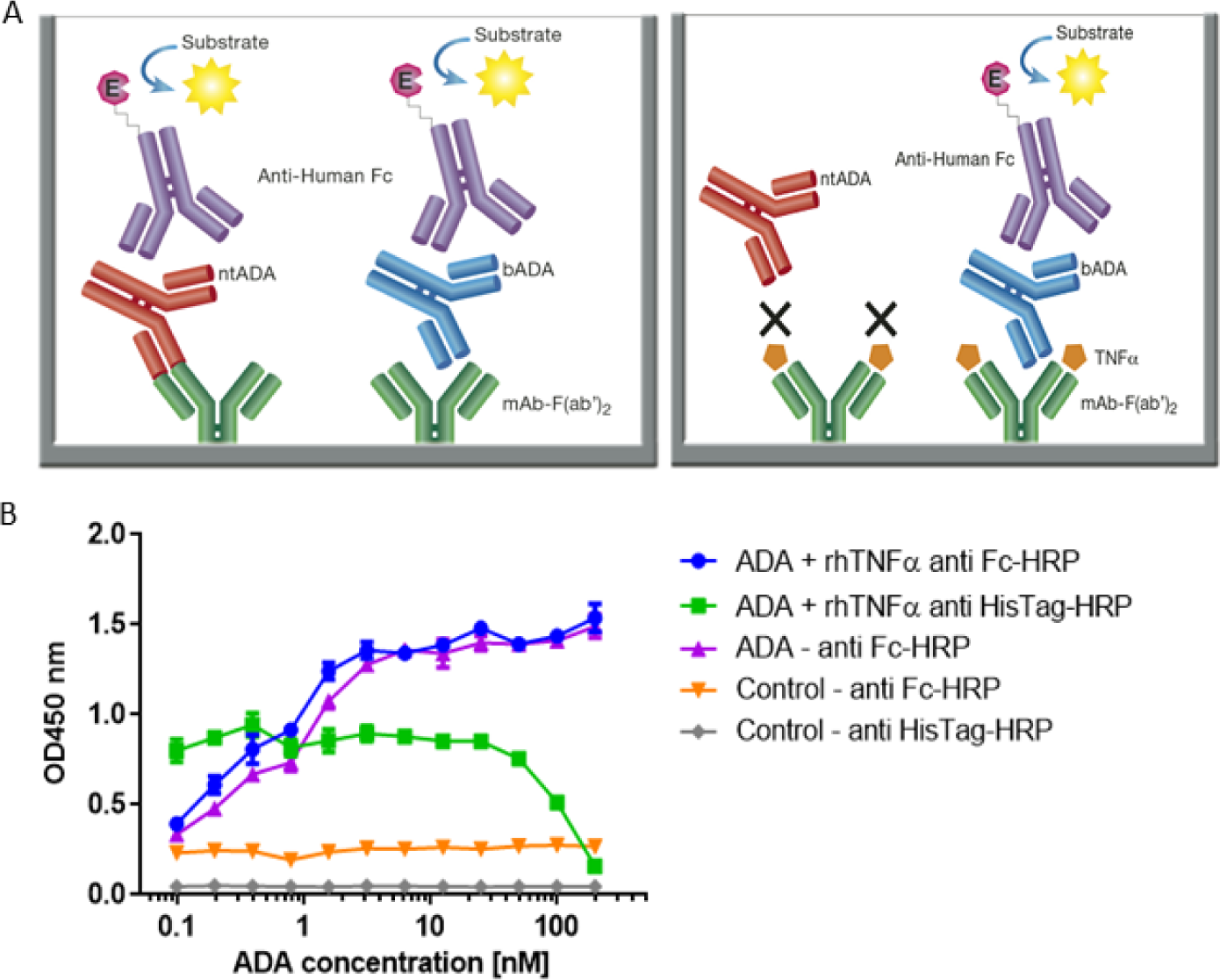
Configuration of the assay for determining the neutralization index of ADA in patient sera and competitive ELISA between ADA and rhTNFα. (A) The newly developed mAb-F(ab’)_2_ based bio-immunoassay configuration (left) and the modified configuration where mAb-F(ab’)_2_ binding site is blocked by saturating the assay with rhTNFα (right). (B) Competitive effect of rhTNFα on ADA binding to IFX-F(ab’)_2_. ELISA plate was coated with 5μg/ml of IFX-F(ab’)_2_. ADA standard was diluted 3-fold in blocking solution supplemented with 5nM rhTNFα. ADA diluted 3-fold in blocking solution without the presence of rhTNFα served as a control.

First, we evaluated the ability of rhTNFα to inhibit the binding of ADA to the coated IFX-F(ab’)_2_ by setting up a competitive ELISA where a series of ADA standard concentrations were incubated with a series of fixed rhTNFα concentrations (data no shown). We observed a competitive effect while rhTNFα was fixed at the concentration of 5nM (Figure 4B). This step was important as it enabled us to determine the ADA equimolar concentration of rhTNFα to be used that will fully occupy the IFX-F(ab’)_2_ binding site and will prevent the binding of *nt*ADA to the coated (and blocked) IFX-F(ab’)_2_. We monitored the presence of rhTNFα using an HRP-conjugated anti-His tag antibody and observed that if we aim to completely block ADA it is required to use equimolar concentration of rhTNFα that is corresponds to the highest concentration of ADA in the assay (200nm).

In practice, IFX-F(ab’)_2_ binding site was blocked with rhTNFα by prior incubation of serum with the coated IFX-F(ab’)_2_ hence, the differential signal w/ and w/o the presence of rhTNFα represent the portion of ADA that could not bind the IFX-F(ab’)_2_ binding site thus, reflects the neutralization capacity (hereby named neutralization index) of the ADA in the tested serum. Using this assay, we evaluated the neutralization index of the 46 ADA positive sera from patients treated with IFX and 7 ADA positive sera from patients treated with ADL. In sera from patients treated with IFX, we noticed that there are two main neutralization index patterns: those with high differential signal (Figure 5A) and low differential signal (Figure 5B). More interestingly, we found that patients that were stratified as AHLC^(+)^ have a significantly higher neutralization index compared to those that belong to the AHLC^(-)^ group (Figure 5C). This suggests that there is a preferential usage of the λ light chain in *nt*ADA as the AHLC^(+)^ group is *a priori* defined by the presence of ADA comprising the λ light chains. All sera from patients treated with ADL (n=7) were subjected to modified bio-immunoassay and demonstrated high neutralization indexes (Figure S2)

**Figure 5:**
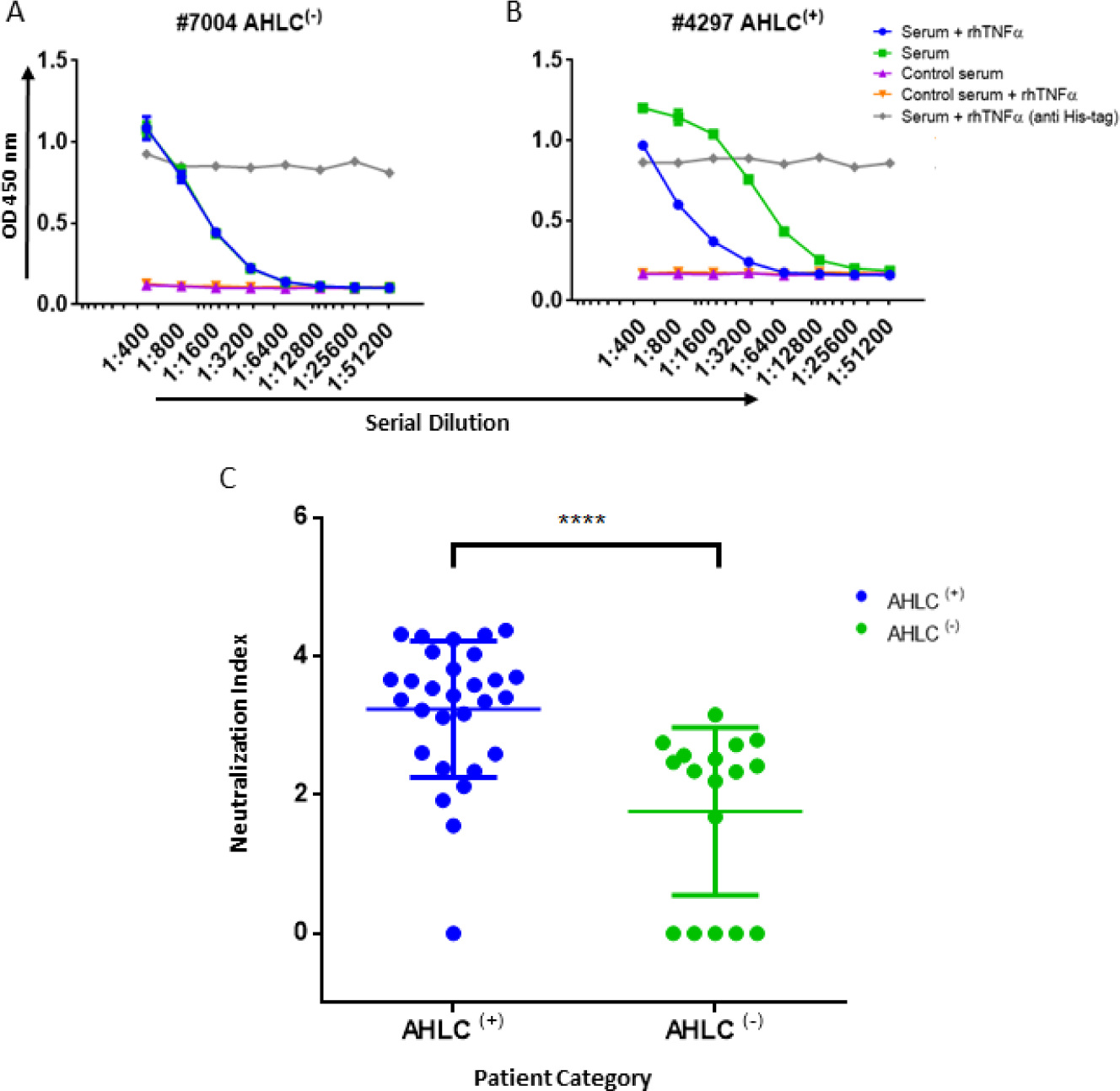
Neutralization index ELISA. (A) Graph representing the ELISA results obtained utilizing the neutralization assay on serum that was designated as AHLC^(-)^ and (B) AHLC^(+)^. In both (A) and (B) the effect of soluble TNFα on ADA detection was evaluated and neutralization index was determined. (C) Scatter plot consolidating the neutralization index obtained by applying the immunoassay on sera from 46 ADA positive patients treated with IFX (\*\*\*\**P* < 0.0001, Mann-Whitney *U* test). For A–C, averages were calculated as mean, with error bars indicating s.d.

### IFX infusion induces a vaccine like immune response

To further investigate the molecular landscape of ADA we explored the dynamics of the B cell response following mAb administration. When investigating well-controlled clinical scenarios such as samples obtained from post-vaccinated individuals, it is convenient to isolate the antigen-specific B cell as they peak at a defined time window (23, 32). However, the characteristics of the humoral response and ADA encoding B cell dynamics following mAb administration is unknown. Our working hypothesis assumed that the immune response following mAb administration is a vaccine-like response thus; we expected to observe a wave of PB peaking several days after IFX infusion. It was previously demonstrated that boost vaccines induce a strong proliferation of PBs and mBCs that can be detected in the blood circulation several days after the boost (33, 34). To test if IFX administration induces a vaccine like response, we collected blood samples from a patient that was found to be positive to ADA at two time points: prior to IFX infusion (D0) and 10 days after IFX infusion (D10). The second time point (D10) was determined in order to capture an enriched population of antigen-specific PB as well as mBC that enable the establishment of a donor-specific V_H_ database for the proteomic interpretation of peptides derived from ADA.

Peripheral blood mononuclear cells (PBMCs) were sorted by FACS and the frequency of PB (CD3^−^CD19^+^CD20^-^CD27^++^CD38^++^) and mBC (CD3^-^CD19^+^CD20^+^CD27^+^) subsets were determined. We identified a 13-fold increase in the frequency of PB at D10 and no increase in the mBC compartment. The PB data suggests that the B cell dynamics following IFX infusion exhibits vaccine-like characteristics in accordance with our working hypothesis (Table 2, Figure S3).

**Table 2:**
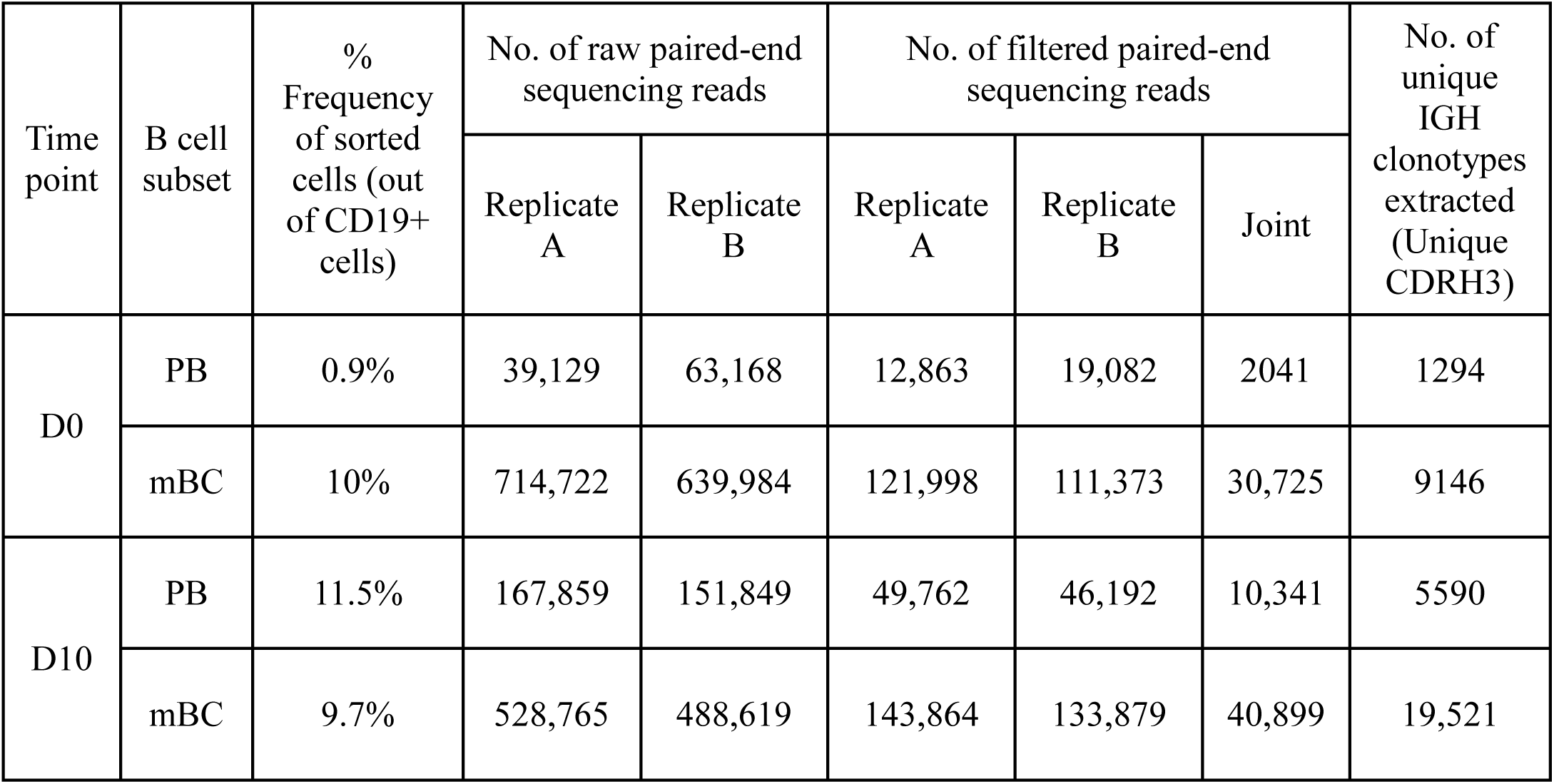
B cell frequency of a patient treated with IFX.

| Time point | B cell subset | % Frequency of sorted cells (out of CD19+ cells) | No. of raw paired-end sequencing reads |  | No. of filtered paired-end sequencing reads |  |  | No. of unique IGH clonotypes extracted (Unique CDRH3) |
| --- | --- | --- | --- | --- | --- | --- | --- | --- |
|  |  |  | Replicate A | Replicate B | Replicate A | Replicate B | Joint |  |
| D0 | PB | 0.9% | 39,129 | 63,168 | 12,863 | 19,082 | 2041 | 1294 |
|  | mBC | 10% | 714,722 | 639,984 | 121,998 | 111,373 | 30,725 | 9146 |
| D10 | PB | 11.5% | 167,859 | 151,849 | 49,762 | 46,192 | 10,341 | 5590 |
|  | mBC | 9.7% | 528,765 | 488,619 | 143,864 | 133,879 | 40,899 | 19,521 |

### Antibody repertoire of ADA encoding B-Cells

The waves of PB following challenge is enriched with antigen-specific B cells (23, 32, 35). Based on this, a major fraction of PB at D10 post mAb infusion is expected to comprise B cell clones responding to the current antigen challenge. Thus, the repertoire of B cells at two time points (pre- and post-infusion) is predicted to represent the overall differences in the ongoing ADA encoding B cell response.

This diversity of antibodies is accomplished by several unique molecular mechanisms, including chromosomal V(D)J rearrangement, somatic hypermutations (SHM) and class switch recombination (25), processes that are mediated by recombination-activating gene (RAG) and activation-induced cytidine deaminase (AID), respectively. The AID enzyme functions mainly in secondary lymph nodes named germinal centers. Next-generation sequencing (NGS) of the antibody variable regions (V-genes) coupled with advanced bioinformatics tools provides the means to elucidate the antigen-specific antibody repertoire’s immense diversity (36). To deep sequences antibodies’ V-genes, recovered RNA from sorted PB and mBC was used as the template for first-strand cDNA synthesis, followed by PCR amplification steps to produce barcoded amplicons of the V-genes of the antibody heavy chains (V_H_) as described previously (24). While NGS of antibodies is a powerful tool for immune repertoire analysis, relatively high rates of errors accumulate during the experimental procedure. To overcome this challenge, we generated duplicates of the antibody V-gene amplicons and sequenced them using the Ilumina MiSeq platform (2×300bp). The resultant V_H_ sequences were processed using our recently reported ASAP webserver that was specifically developed to analyze NGS of antibody V-gene sequences derived from replicates (25).

In our analysis, we concentrated on several repertoire measures that collectively provide a molecular level characterization of the ADA: i) V(D)J family usage; ii) CDR3 length distribution; iii) SHM levels, and, iv) isotype distribution. Our data revealed several interesting antibody repertoire features that may shed light on the molecular mechanism involved in the formation of ADA.

#### V(D)J gene family usage is stable

Examining the V(D)J family usage is important to determine whether the basal gene frequency is similar to the expected frequency and if the B cell response following IFX infusion drives B cells to exhibit a preferential V(D)J gene usage. Therefore, we examined the frequency of family usage at two time points (D0 and D10), within PB and mBC subsets across isotypes (IgG and IgM). The V(D)J family usage showed no marked difference between the two time points, B cell subsets and isotypes. The frequency of V-gene family usage was also found to have similar frequency profile as previously described (37, 38). For example, the V-gene family frequency showed that the V3, V4 and V1 have the most prevalent representation followed by V2, V5 and V6 that had significantly lower frequencies (Figure 6A). The same pattern trends were identified for the D and J family usage.

**Figure 6:**
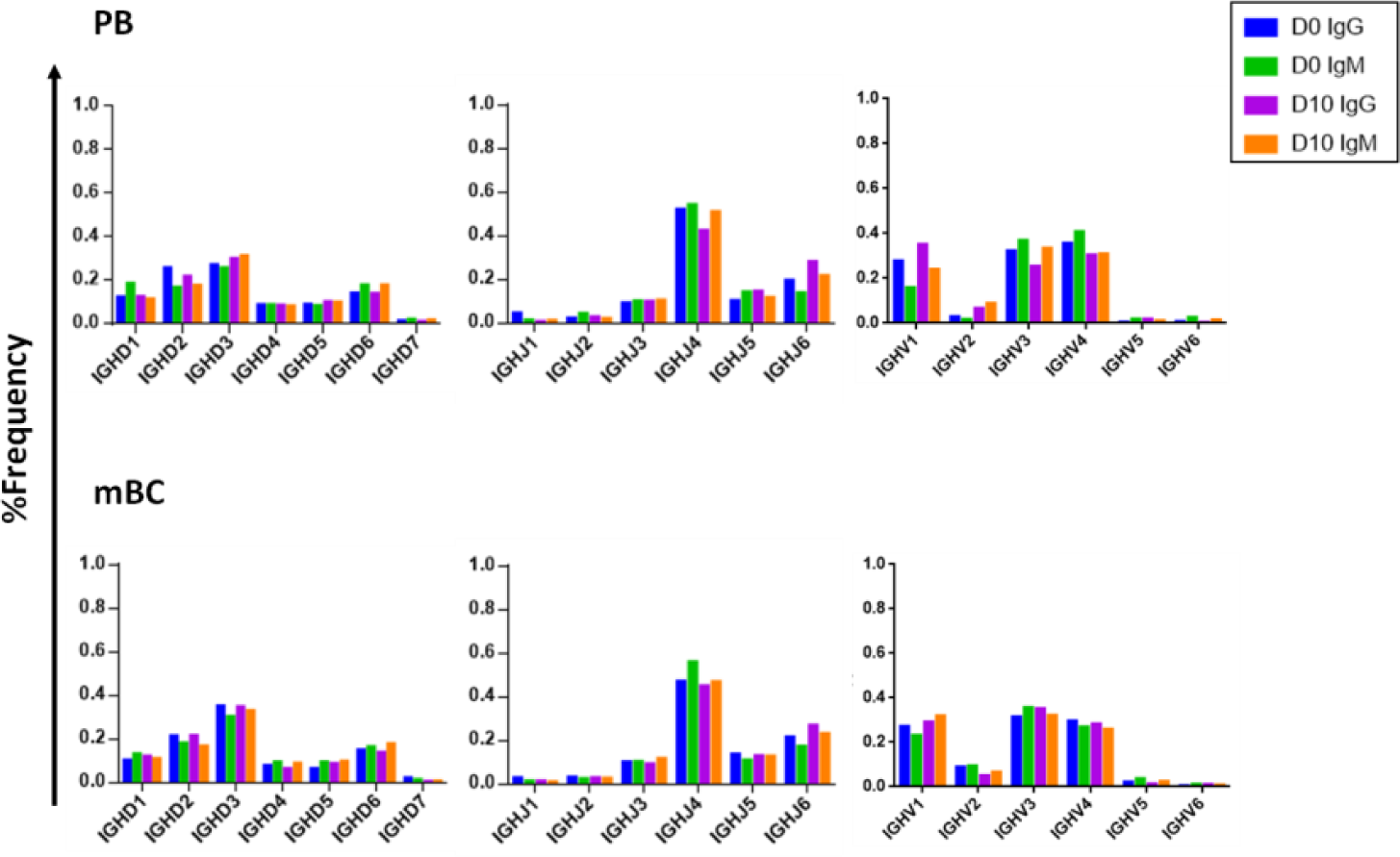
V, D and J family usage in B cell following IFX infusion. mBC and PB from a patient treated with IFX were collected at two time points (D0, D10) and processed for NGS analysis. The V family usage showed no difference between D0 and D10, different B cell subsets and isotypes. The D and J family usage showed no difference between time points.

#### CDRH3 length increases following IFX infusion

Composed of the V(D)J join with its inherent junctional diversity, the CDRH3 specifies the antibody V_H_ clonotype. The V_H_ clonotype is an important immunological concept because it accounts for antibodies that likely originate from a single B-cell lineage and may provide insight on the evolution of the antigen-specific response (39). Here we defined V_H_ clonotype as the group of V_H_ sequences that share germ-line V and J segments and have identical CDRH3 sequences. By examining the length distribution of CDRH3 from PB across isotypes and time point we observed a shift towards longer CDRH3 at D10 (Figure 7). Interestingly, this observation is in contrast to previous studies that reported a decrease in the CDRH3 length post immunization with pneumococcal (40) and hepatitis B vaccines (41) and when comparing antigen experienced B cell to naïve B cells (42).

**Figure 7:**
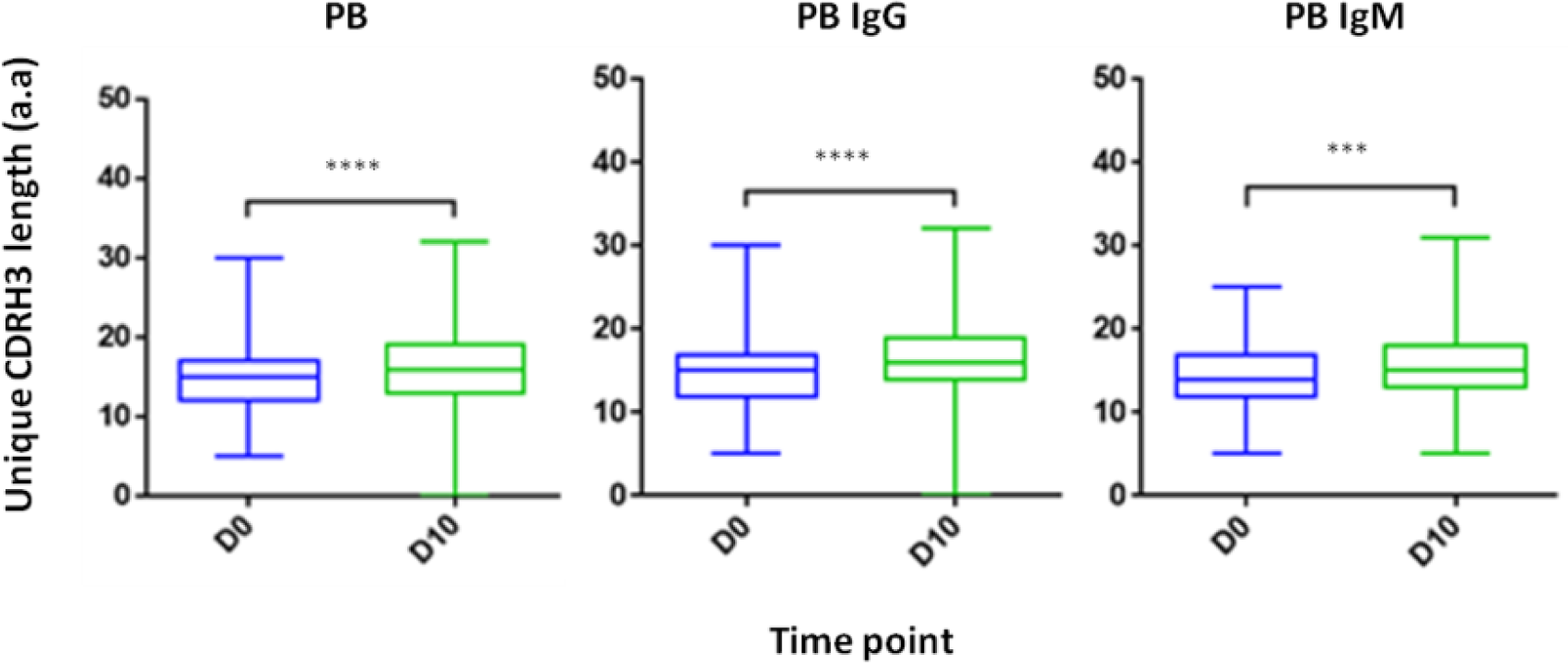
CDRH3 length at two time point and across isotypes. PB from a patient treated with IFX were collected at two time points (D0, D10) and processed for NGS analysis. An increase in antibody CDRH3 length was observed. (\*\*\**P*<0.001, Mann-Whitney *U* test).

#### Somatic hypermutation levels decreases following IFX infusion

Examining the level of SHM following vaccination provides insights regarding the extent of the affinity maturation that antigen-stimulated B cell undergo. It was previously reported that boost vaccination induces a substantial increase of the SHM levels when comparing post- to pre-vaccination (41). Despite the vaccine like response following IFX infusion, we observed in the PB compartment a significant decrease in the SHM levels post-infusion, regardless if the mutations were synonymous and non-synonymous (Figure 8).

**Figure 8:**
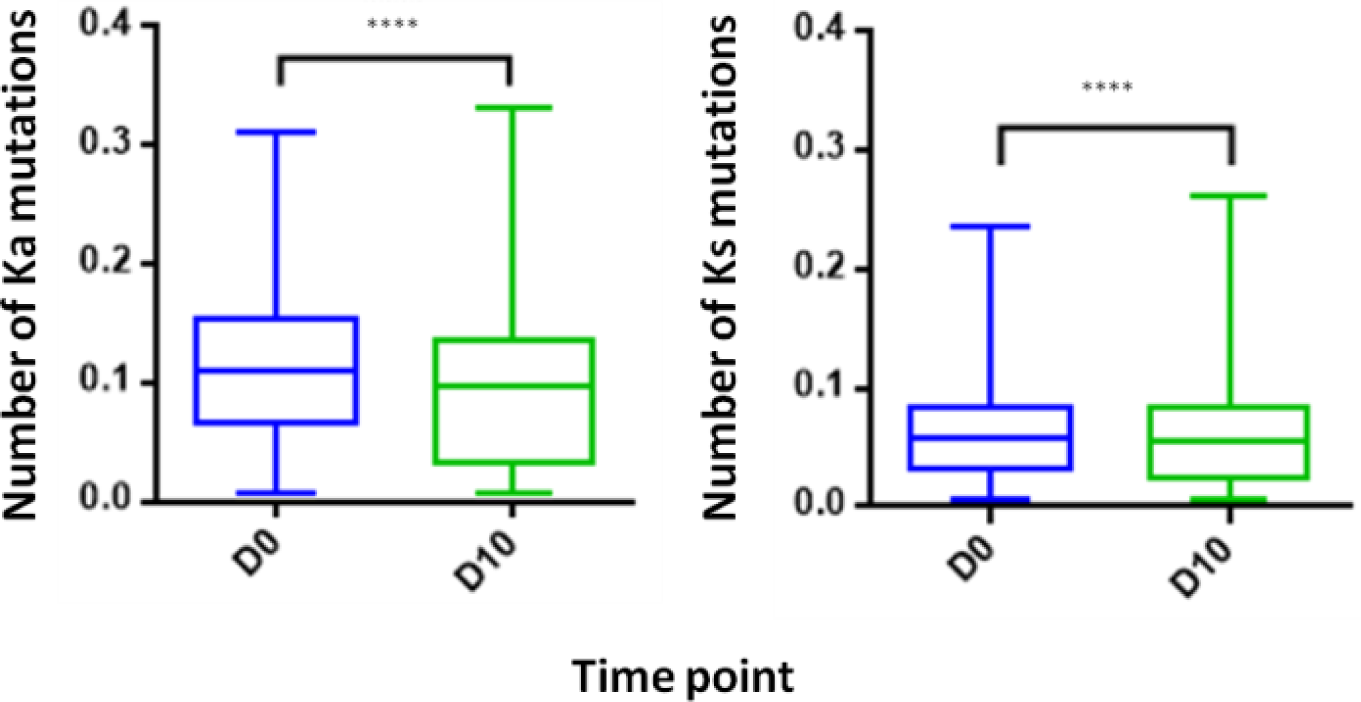
Somatic hyper mutations. PB from a patient treated with IFX were collected at two time points (D0, D10) and processed for NGS analysis. A decrease in the number of Ka mutations (number of non-synonymous mutation per codon) and Ks mutations (number of synonymous mutations per codon) was observed at D10. (\*\*\*\**P*<0.0001, Mann-Whitney *U* test).

### Proteomic analysis of ADA

Analysis of serum antibodies provides a comprehensive profile of the humoral immune response and is complementary to the transcriptomic analysis derived from NGS of the antibody V_H_. Applying an approach that integrates NGS and tandem mass spectrometry (LC-MS/MS) has been shown to provide valuable data regarding the composition of antigen-specific serum antibodies and their relationship to B cells and generates new insights regarding the development of the humoral immune response in disease and following vaccination (23, 32). Here we utilized the previously developed omics approach (26, 39) to elucidate the serum ADA composition following IFX infusion. ADA from 10 ml of serum collected at D0 and D10 were subjected to protein G affinity chromatography and total of 9 mg of recovered IgG was digested by IdeS to remove the Fc regions that may mask the MS/MS signal obtained from low abundant peptides. Following 5 hours of digestion, the reaction mixture was subjected to custom made affinity column where the IFX-F(ab’)_2_ was coupled to agarose beads and served as the antigen to isolate ADA. Recovered 48.57µg polyclonal ADA-F(ab’)_2_ (i.e., IFX-F(ab’)_2_-specific F(ab’)_2_) in the elution fraction and total F(ab’)_2_ (depleted from ADA-F(ab’)_2_) in the flow through fraction were digested with trypsin and injected to high-resolution tandem mass spectrometer analyzer in triplicates. LC-MS/MS raw data files were analyzed using MaxQuant using label free quantitation mode (LFQ) and searched against the custom antibody V-gene database derived from the NGS data of B cells isolated from the same donor. Identified peptide from the interpretation of the proteomic spectra were stratified into three types of peptides: informative peptides (*i*Peptide) that map uniquely to one antibody clonotype in a region that is upstream to the CDRH3, non-informative CDRH3 peptides (*ni*CDRH3) that map to the CDRH3 region of the antibody but do not map uniquely to a single antibody clonotype and informative CDRH3 peptides (*i*CDRH3) that map uniquely to a single antibody clonotype. Summary of identified peptides in LC-MS/MS are shown in Table 3. Beyond the designation as *i*CDRH3 peptides, additional filtration steps were applied including peptides that were present in more than 2 replicates, peptides in elution that show 5xfold frequency than in the flow through. The *i*CDRH3 peptides enabled the identification of 62 unique ADA CDRH3 clonotypes with 205 associated full-length V-gene sequences. The resulting V-gene sequences were analyzed to determine their V(D)J family usage and the B cell subset they are mapped to, based on our NGS data (Figure 9).

**Table 3:** Summary of identified peptides and the corresponding clonotype and antibody somatic variences in the LC-MS/MS spectra. E: elution, FT: flow-through.

|  | Day 0 | Day 10 |
| --- | --- | --- |
| Total peptides | 908 | 3177 |
| Total antibody peptides | 761 | 2805 |
| Total CDRH3 | 42 | 224 |
| Present in ≥2 technical replicates | 30 | 166 |
| Frequency ratio E/FT > 5 | 11 | 81 |
| Nº of clones | 5 | 62 |
| Nº of somatic variances | 35 | 205 |

**Figure 9:**
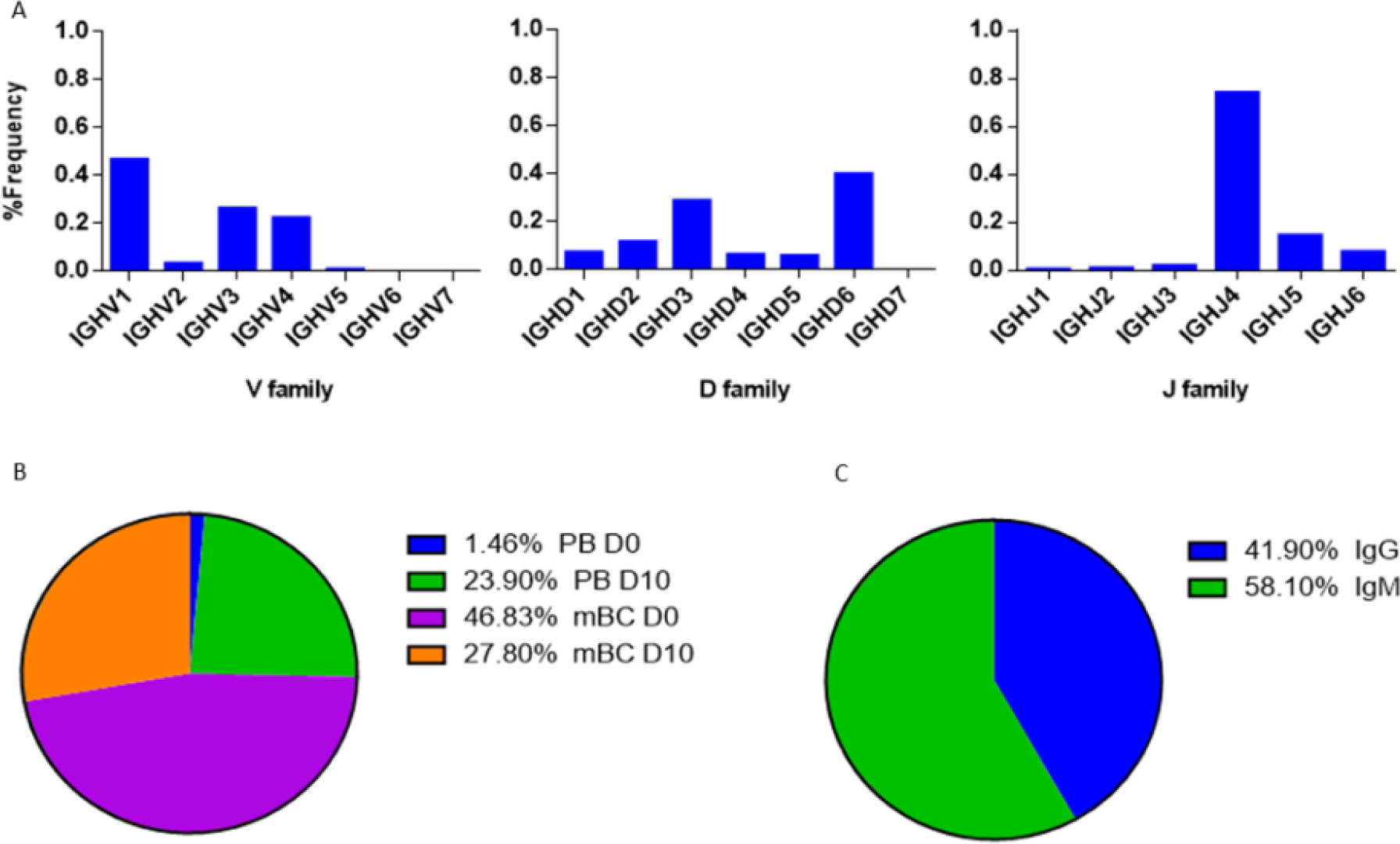
V-gene and circulating antibody repertoire characteristics. (A) The V(D)J family usage of V-gene sequences that were identified by LC-MS/MS. (B) Mapping of V-gene sequences to B cell subsets and (C) isotypes, based on NGS data.

The V(D)J family usage of the antibody variable region sequences that were identified by LC-MS/MS (Figure 9A) showed a similar distribution as observed in the NGS data (Fig. 6A, Fig. 7A). V family frequency analysis showed that the V1, V3, and V4 are the most dominant V families followed by V2 and V5 that had significantly lower frequencies. D family frequency analysis showed that the D6, D3, D2 and D1 have the most prevalent representation, and J family frequency showed that the J4, J5 and J6 have the most prevalent representation.

Next, we examined the distribution of the proteomically identified V-gene sequences to B cell subsets (Figure 9B) and found that the V-genes predominantly map to mBC from D0 (46.83%), followed by mBC from D10 (27.8%). Moreover, we found that 23.9% of V-genes map to D10 PB. Based on the dynamics of antibodies in serum (32), the majority of antibodies produced following a boost challenge are the product of pre-existing mBC cells that were re-activated following drug infusion, much like a response to a vaccine boost (23).

As mentioned above, flow cytometry of B cells following IFX administration allowed us to identify a substantial increase in the frequency of PB at D10, which suggests that the B cell dynamics following IFX infusion exhibits vaccine-like characteristics. Therefore, we expected to find a majority of V-gene sequences mapping to IgG^+^ B cells that underwent class switch recombination in the germinal center. Surprisingly, the majority of proteomically identified serum antibodies were mapped to IgM^+^ B cells (Figure 9C).

Next, we aimed to provide support to the observation that *nt*ADA preferably use the λ light chain. By quantifying the accumulative intensities of peptides derived from the constant region of both κ and λ light chains, we calculated the ratio of κ:λ light chain in the elution fraction which comprise both the *nt*ADA and *b*ADA (ADA-IgG), and in the flow through fraction that represent ADA-depleted IgG (dep-IgG). The expected κ/λ ratio of IgG in human serum is 2 (66% κ and 33% λ). Indeed, proteomic analysis of the dep-IgG (D0 and D10), resulted in an average κ/λ ratio of 2.1. The same analysis of the ADA-IgG showed a significant shift of the κ/λ ratio to 1.19. The proteomic analysis was carried out on samples from patient that exhibited a high neutralization index (have high levels of *nt*ADA) and designated ADA^+^ using AHLC (detects only ADA with λ light chain). Brought together, this further suggests that *nt*ADA contribute to shift in the k:λ light chain ratio.

## Discussion

The use of therapeutic mAbs in treating a wide range of diseases and disorders is growing exponentially. Nonetheless, a major shortcoming of their use is the development of ADA in patients receiving the mAb. Advances in mAb engineering have enabled the development of fully human mAbs with reduced immunogenicity without abolishing it completely. Thus a mAb administered to a patient can still induce an immune sensitization as reflected by the production of ADA, which is associated with low trough drug levels and can mediate loss of clinical response to the drug (20).

The precise mechanism underlying ADA production is unknown, and many questions related to its development remain unaddressed, including determining precise concentrations of ADA in serum, which portion of the ADA exhibits neutralizing capacity, the immune pathway governing the production of ADA, and ultimately, the molecular composition of ADA at the sequence level. To address these questions, we chose the chimeric TNFα antagonist IFX as the model system. First, we aimed to quantify the ADA level in patient sera. Many methods were previously reported to evaluate serum ADA levels. These assays include radio-immunoassays (43), Biotin-drug Extraction with Acid Dissociation (BEAD) (44), Precipitation and acid dissociation (PANDA) (45), affinity capture elution ELISA (ACE) (46), and Homogenous Mobility Shift Assay (HMSA) (47). While these assays are not limited to the λ light chain detection like the AHLC assay, they provide mostly qualitative measures to assist physicians in deciding the most appropriate intervention when treating patients, and many (if not all) studies underestimated actual ADA levels (19). These assays also lack a standardization methodology that can enable the comparison of ADA levels across health centers.

To provide quantitative measures describing the molecular landscape of ADA, we first developed a bio-immunoassay that would allow quantify ADA levels based on the F(ab′)_2_ region of the mAb because previous reports indicated that the ADA generated from mAb administration are mostly anti-idiotypic (21). Indeed, the bio-immunoassay demonstrated higher sensitivity compared with the AHLC assay used initially to detect ADA and was able to detect ADA when the AHLC assay could not. Leveraging its improved sensitivity compared to the AHLC assay, we applied our proprietary assay on sera from 54 patients treated with IFX and found that patients designated as AHLC^(+)^ showed significantly higher levels of ADA (mean: 264 μg/ml) compared to the AHLC^(-)^ group (mean: 59.64 μg/ml). These results support the clinical use of AHLC assay because overall, patients were correctly stratified leading to clinical decision-making that was based on a valid indicative assay. Notwithstanding, the applicability of the AHLC assay, the newly developed F(ab′)_2_-based bio-immunoassay demonstrated that ADA levels can reach extreme concentrations that were not detected using the AHLC assay.

Some patients who develop ADA in response to IFX present a prolonged remission with maintenance therapy despite repeated indications of high ADA and low IFX trough levels (20). The mechanism of action of these ADA has significant influence on drug efficacy. For example, *b*ADA are most likely to enhance the clearance of a drug whereas *nt*ADA will prevent a drug from binding to its target. Hence, it is important to differentiate between *b*ADA and *nt*ADA, or in other words, a need exists to identify sera with high levels of *nt*ADA that may predict the likelihood of a patient losing a favorable response to an administrated mAb. To achieve this, we further revised our bio-immunoassay to qualitatively measure the neutralization index of ADA in the serum of patients treated with IFX. Of note, as the neutralization index is a qualitative and not a quantitative index, some patients may exhibit relatively low ADA levels and high neutralization index. Using this assay on sera from the 46 ADA positive patients, revealed that patients who tested positive utilizing the AHLC assay, exhibit a significantly higher neutralization index than patients tested negatively for it (i.e., AHLC^(-)^). Noteworthy, the AHLC assay is based on the anti-λ light chain antibody at the detection stage, suggesting that sera with high neutralization index comprise ADA that preferably use the λ light chain (either *b*ADA or *nt*ADA). This phenomenon received additional support from our proteomic analysis in which we compared the changes in the ratio between peptides derived from κ and λ constant light chains from ADA-IgG pool and peptide derived from depleted ADA IgG polyclonal pool (dep-IgG). This analysis demonstrated that the κ/λ ratio in the total IgG compartment is as expected and is decreased in the mAb-specific compartment (κ/λ ratio 2.1 and 1.19 for dep-IgG and ADA-IgG, respectively). The preferential use of the λ light chain in neutralizing antibodies has been previously reported (21, 48), however, the authors of those studies did not provide an explanation beyond the structural adaptability of the light chain toward the target. The relevance of the reported cases showing λ chain bias is not clear. Similar phenomena was reported in B-1 sub-population, unlike follicular B cells, B-1 cells exhibit an increased frequency of lambda light chains (49). The recurrence of BCRs with the enrichment of λ light chain has been considered to result from strong antigen-dependent selection of the B-1 cell repertoire (50).

Repetitive administration of mAbs may induce a strong humoral response manifested in the production of ADA. We hypothesized that mAb administration is similar to the response that occurs following a boost vaccine. Others and we have demonstrated that boost vaccines induce a strong proliferation of PB that can be detected in blood circulation several days after the boost. The “wave” of B cells after the boost vaccine are dominated by antigen-specific B cell (34) thus, repertoire analysis of these cells can provide invaluable data about the antigen-specific antibody repertoires. Utilizing flow cytometry showed an order of magnitude increase in PB compartment 10 days after IFX infusion, suggesting that the immune response following IFX administration is indeed similar to a vaccine response. To the best of our knowledge, this is the first report to identify a vaccine like response following therapeutic mAb administration.

Next, we aimed to provide a comprehensive repertoire profile of the B cells induced after mAb administration. To achieve this, we applied an “omics” approach as previously described (23, 26, 39) that is based on the integration of NGS of the V-genes and proteomic analysis of serum ADA. NGS of V-genes revealed no bias in the V(D)J usage across isotypes, cell types, and time point. These data suggest that the original repertoire that existed before mAb administration and antigen-specific repertoire induced by IFX administration is formed by random recombination processes without preferential use of any particular V(D)J segment. Comparative repertoire analysis of the V-genes between time points (before and after IFX administration) revealed that post-IFX administration, PB exhibit longer CDRH3 and lower SHM rates. Although the B cell dynamics after mAb administration are similar to those that occur after a boost vaccine, the repertoire measures show a different profile. It was previously reported that the antibodies generated after a boost vaccine exhibit shorter CDRH3, high SHM (40-42).

To explain these data we revisited two reports: the first describes how the immune response in TNFα-deficient mice was “diverted” to the marginal zone instead of to the germinal center (51) and the characteristics of the immune response in the marginal zone is directly affected by low levels of the AID that in turn is reflected in lower SHM rate. The second reported a skewed λ chain usage in B-1 cells (49). Based on these reports we propose a mechanistic model according to which administration of TNFα antagonist blocks the TNFα on one hand and induces a vaccine-like response on the other. Due to the TNFα blockade, immune response of B cells occurs extra follicular where AID is downregulated, thus the encoded ADA exhibit lower SHM rates. Moreover, the data suggests that the immune response following mAb administration may be a T cell independent (TI) response which is governed by the B-1 cell linage with the characteristics mentioned of an increased usage of λ light chains and little to non-evidence for SHM (49, 52).

Another possible mechanism that should be further explored is the strong TI immune response in the marginal zone that is also induced by a drug/ADA immune-complex (IC). It was previously suggested that many of the immune-mediated adverse effects attributed to ADA require the formation of an IC intermediate that can have a variety of downstream effects (6, 53).In the context of the system we investigated, administration of a TNFα antagonist will divert the immune response extra follicular either by TNFα blockade or by the formation of an IC carrying multiple mAbs that can induce the cross-linking of cognate BCR. The BCR of ADA-encoding B cells will undergo co-clustering leading to their activation in the TI pathway.

Of note, insights from this study are restricted to the immune response following treatment with TNFα antagonists, as it is specifically affected by the drug’s mechanism of action. First, TNFα is a trimer that has the propensity to form immunocomplexes with the drug. Second, blocking TNFα “simulates” a scenario that was observed in TNFα knockout mice. Combined, these attributes contribute to the specific nature of the immune response which is suggested to be diverted to the extra follicular, TI immune response. Moreover, the deep analysis data was obtained from one patient. However, this patient exhibited a set of attributes including high ADA level, high neutralization index, low trough level and lack of immunosuppressant treatments. These attributes enabled to generate insights that are directly relevant to the drug administration.

In our study we examined molecular aspects related to the formation of ADA. To the best of our knowledge, this is the first report describing ADA repertoire that resulted in insights about a possible mechanism of ADA formation. Further work will be needed to elucidate additional phenotypic markers of the B cells induced by mAb administration and the role of IC in the activation of the B cell. Moreover, the mechanism described here covers the response to a TNFα-antagonist, and by using the same omics approaches, it will be highly informative to study the B cell response following treatment with other mAbs that induce ADA formation. We envision that high throughput data obtained from such studies can facilitate our understanding on why and how, mAb administration generates ADA and eventually may contribute guidelines for engineering therapeutic mAbs with reduced immunogenicity.

## General

We are grateful to George Georgiou for assisting with the LC-MS/MS measurements at UT Austin, for Ulrich von Pawel-Rammingen from the Department of Molecular Biology, Umea University who kindly donated the plasmid with the gene encoding the IdeS.

This manuscript has been released as a Pre-Print at BioRxiv. DOI: https://doi.org/10.1101/509489

## Author contributions

A.V.M, S.B.H, I.B. and Y.W. conceived the research. A.V.M. S.R. and Y.W. designed the experiments, A.V.M., S.R., M.Y., E.F. and Y.D. performed experiments, A.V.M., S.R., S.B.H, M.Y., E.F., B.U. and U.K. collected and processed clinical samples, A.V.M, A.K. and Y.W. carried out data analysis, A.V.M. and Y.W. wrote the manuscript.

