## Supplemental file for "Molecular Landscape of Anti-Drug Antibodies Reveals the Mechanism of the Immune Response Following Treatment with TNFα Antagonists"

### Supplementary material

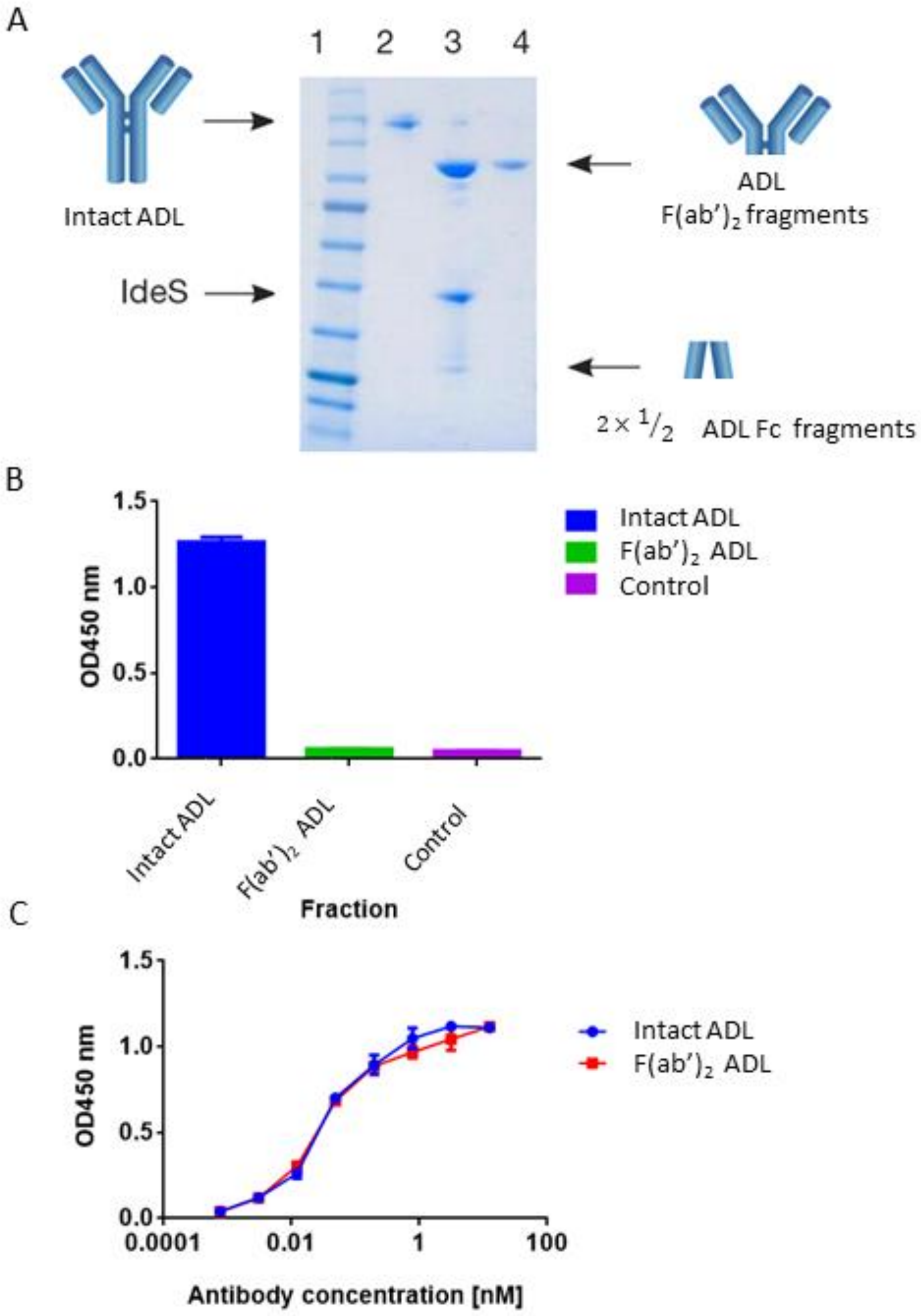


**Figure S1**: **ADL digestion and ADL-F(ab')$_2$ purification**. (A) SDS-PAGE analysis of intact ADL (lane 2), following IdeS digestion (lane 3) and purified ADL-F(ab')$_2$ following a 2-step affinity chromatography purification including protein A and kappaSelect columns (lane 4). (B) Presence of ADL-Fc and intact ADL traces was measured by direct ELISA were intact ADL and purified ADL-F(ab')$_2$ were compared to a control antigen (streptavidin) as coating agents followed by direct incubation with an anti-Fc HRP conjugate at the detection phase. (C) The functionality of the recovered ADL-F(ab')$_2$ was confirmed by ELISA and compared to intact ADL. The ELISA setup included TNF$\alpha$ as the coating agent and anti-$\kappa$ HRP conjugate at the detection phase. For panel B–C, triplicate averages were calculated as mean, with error bars indicating s.d.

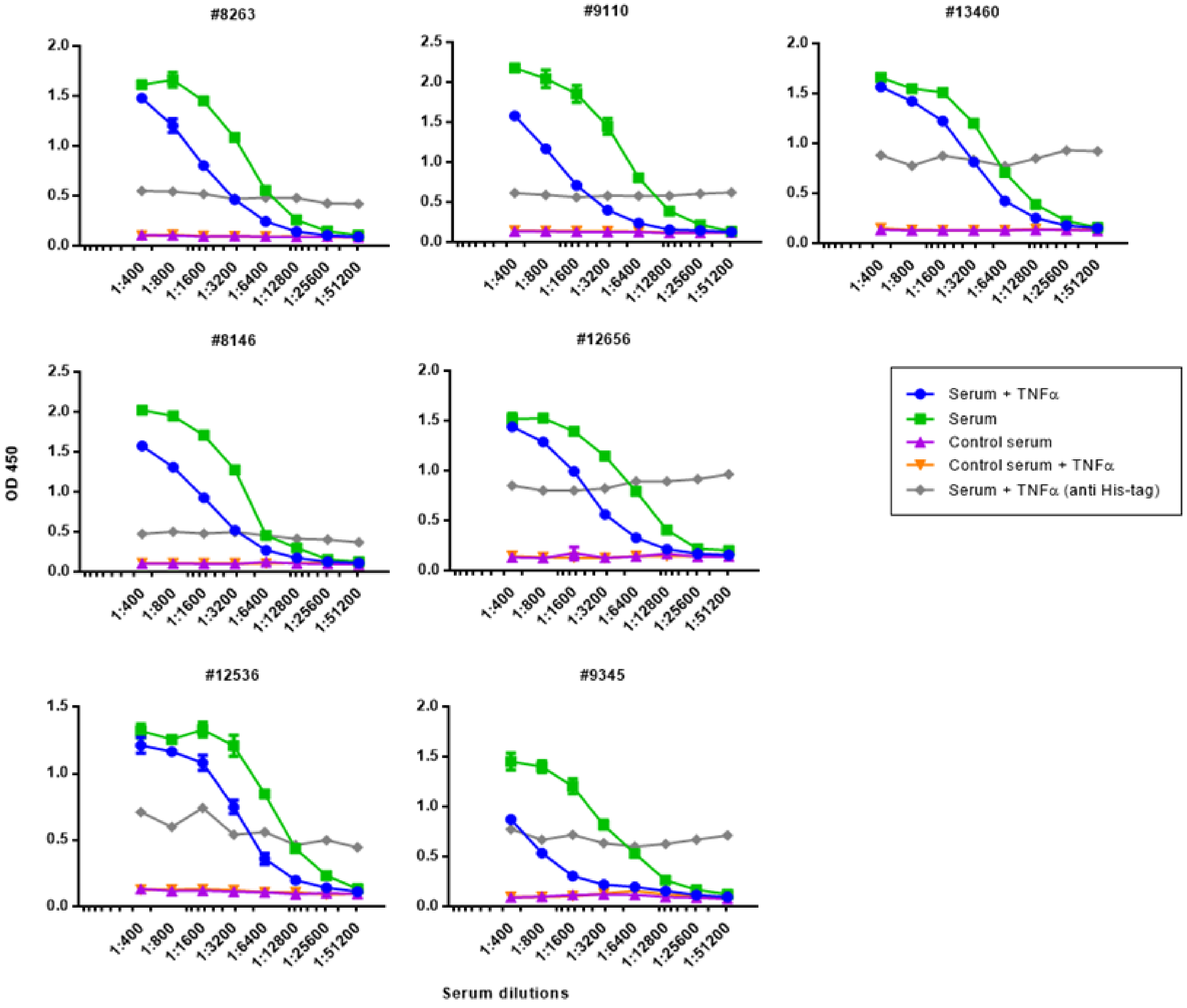


**Figure S2: Determination of ADA neutralization index in patients treated with ADL.** The reduction of the signal obtained when *nt*ADA are present in serum was indicative for sera with high neutralizing capacity.

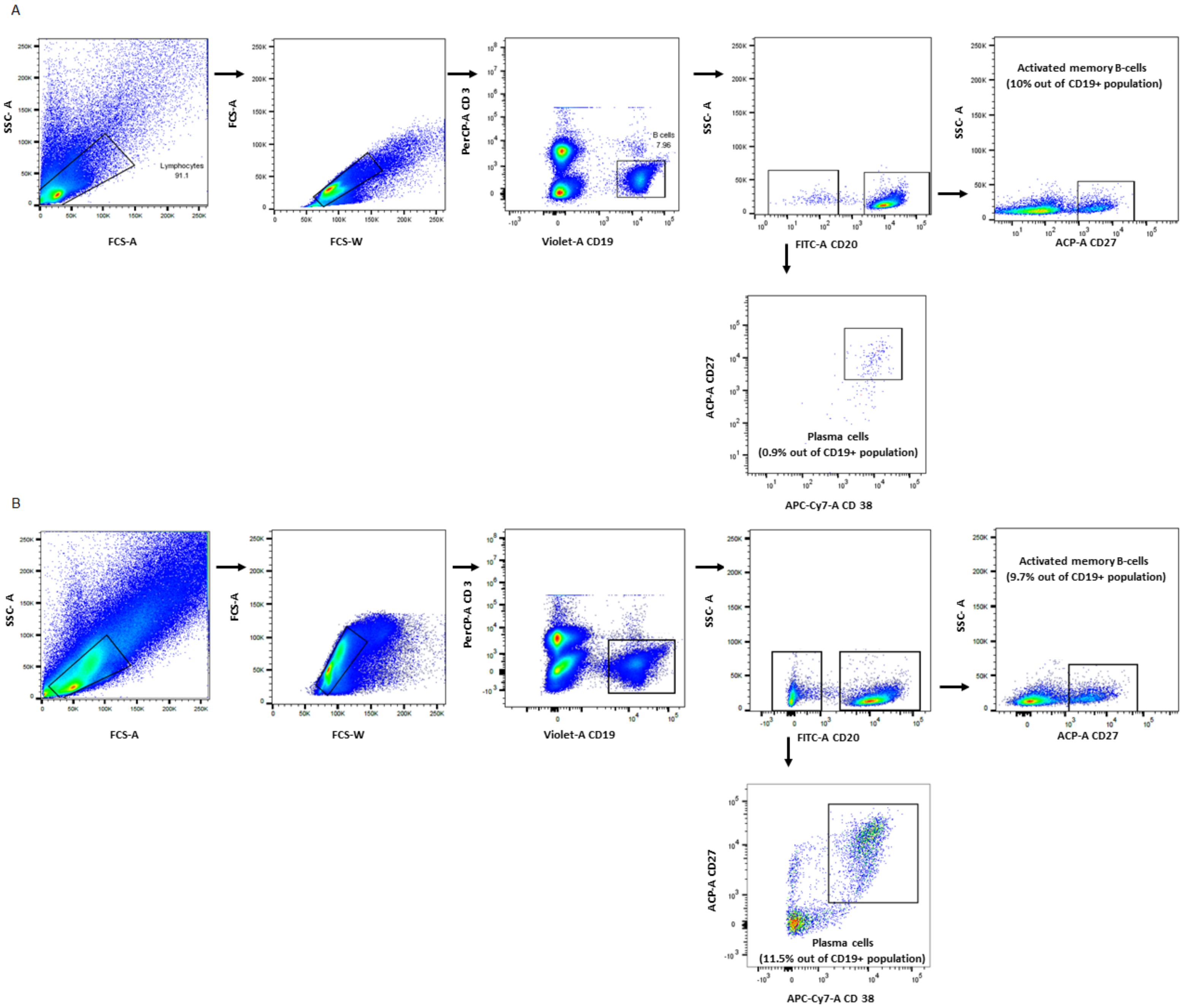


**Figure S3:** FACS of PB and mBC isolated from peripheral blood of IBD patient treated with IFX. (A) Gating on plasmablasts ($CD3^-CD19^+CD20^-CD27^{high}CD38^{high}$) and mBC ($CD3^-CD19^+CD20^+CD27^+$) mBC at D0. Approximately 0.9% of all B cells stained as PB, and 10% as mBC. (B) FACS of $CD3^-CD19^+CD20^-CD27^{high}CD38^{high}$ plasmablasts and $CD3^-CD19^+CD20^+$ mBC at D10. Approximately 11.5% of all B cells stained as PB, and 9.7% as mBC.

**Table S1:** List of all primers used for amplification of $V_H$ genes. fw: forward, rev: reverse, UAd: universal adapter, Idx: index, RC: reverse complement.

**PCR1**

| **IgH extansion forward** | **Extension + VH 5' specific region** |
|---|---|
| VH1-fw | CCCTCCTTTAATTCCC CAGGTCCAGCTKGTRCAGTCTGG |
| VH157-fw | CCCTCCTTTAATTCCC CAGGTGCAGCTGGTGSARTCTGG |
| VH3N-fw | CCCTCCTTTAATTCCC TCAACACAACGGTTCCCAGTTA |
| VH2-fw | CCCTCCTTTAATTCCC CAGRTCACCTTGAAGGAGTCTG |
| VH3-fw | CCCTCCTTTAATTCCC GAGGTGCAGCTGKTGGAGWCY |
| VH6-fw | CCCTCCTTTAATTCCC CAGGTACAGCTGCAGCAGTCA |
| VH4-fw | CCCTCCTTTAATTCCC CAGGTGCAGCTGCAGGAGTCS |
| VH4-DP63-fw | CCCTCCTTTAATTCCC CAGGTGCAGCTACAGCAGTGGG |

| **IgH extansion reverse** | **Extension (RC) + Ig constant specific region (RC)** |
|---|---|
| IgM-human-rev | GAGGAGAGAGAGAGAG GGTTGGGGCGGATGCACTCC |
| IgG-human-rev | GAGGAGAGAGAGAGAG SGATGGGCCCTTGGTGGARGC |
| IgA-human-rev | GAGGAGAGAGAGAGAG GGCTCCTGGGGGAAGAAGCC |

**PCR2**

| **IgALL universal forward** | **TruSeq universal adapter + Diversity region + Extension** |
|---|---|
| IgALL-UAd-fw | AATGATACGGCGACCACCGAGATCTACACTCTTTCCCTACACGACGCTCTTCCGATCT NNNN CCCTCCTTTAATTCCC |

| **IgALL index reverse** | **TruSeq universal adapter (RC) + Diversity region + Extension (RC)** |
|---|---|
| PE-Idx1-rev | CAAGCAGAAGACGGCATACGAGATCGTGATGTGACTGGAGTTCAGACGTGTGCTCTTCCGATCT NNNN GAGGAGAGAGAGAGAG |
| PE-Idx2-rev | CAAGCAGAAGACGGCATACGAGATACATCGGTGACTGGAGTTCAGACGTGTGCTCTTCCGATCT NNNN GAGGAGAGAGAGAGAG |
| PE-Idx3-rev | CAAGCAGAAGACGGCATACGAGATGCCTAAGTGACTGGAGTTCAGACGTGTGCTCTTCCGATCT NNNN GAGGAGAGAGAGAGAG |
| PE-Idx4-rev | CAAGCAGAAGACGGCATACGAGATTGGTCAGTGACTGGAGTTCAGACGTGTGCTCTTCCGATCT NNNN GAGGAGAGAGAGAGAG |
| PE-Idx5-rev | CAAGCAGAAGACGGCATACGAGATCACTGTGTGACTGGAGTTCAGACGTGTGCTCTTCCGATCT NNNN GAGGAGAGAGAGAGAG |
| PE-Idx6-rev | CAAGCAGAAGACGGCATACGAGATATTGGCGTGACTGGAGTTCAGACGTGTGCTCTTCCGATCT NNNN GAGGAGAGAGAGAGAG |
| PE-Idx7-rev | CAAGCAGAAGACGGCATACGAGATGATCTGGTGACTGGAGTTCAGACGTGTGCTCTTCCGATCT NNNN GAGGAGAGAGAGAGAG |
| PE-Idx8-rev | CAAGCAGAAGACGGCATACGAGATTCAAGTGTGACTGGAGTTCAGACGTGTGCTCTTCCGATCT NNNN GAGGAGAGAGAGAGAG |
| PE-Idx9-rev | CAAGCAGAAGACGGCATACGAGATCTGATCGTGACTGGAGTTCAGACGTGTGCTCTTCCGATCT NNNN GAGGAGAGAGAGAGAG |
| PE-Idx10-rev | CAAGCAGAAGACGGCATACGAGATAAGCTAGTGACTGGAGTTCAGACGTGTGCTCTTCCGATCT NNNN GAGGAGAGAGAGAGAG |
